# Epistasis between promoter activity and coding mutations shapes gene evolvability

**DOI:** 10.1101/2022.06.07.495002

**Authors:** Angel F. Cisneros, Isabelle Gagnon-Arsenault, Alexandre K. Dubé, Philippe C. Després, Pradum Kumar, Kiana Lafontaine, Joelle N. Pelletier, Christian R. Landry

## Abstract

The evolution of protein-coding genes proceeds as mutations act on two main dimensions: regulation of transcription level and the coding sequence. The extent and impact of the connection between these two dimensions are largely unknown because they have generally been studied independently. By measuring the fitness effects of all possible mutations on a protein complex at various levels of promoter activity, we show that expression at the optimal level for the WT protein masks the effects of both deleterious and beneficial coding mutations. Mutants that are deleterious at low expression but masked at optimal expression are slightly destabilizing for individual subunits and binding interfaces. Coding mutations that increase protein abundance are beneficial at low expression but incur a fitness cost at high expression. We thereby demonstrate that expression level can dictate which coding mutations are beneficial or deleterious.

## Main Text

Mutations drive evolution through multiple effects, from altering gene expression to modifying the stability, assembly and function of proteins, which ultimately affect fitness. Therefore, measuring the fitness effects of mutations and identifying the underlying molecular mechanisms has been a long-standing goal in evolutionary biology (*1*, *2*). Due to epistasis, the effects of new mutations are dependent on prior mutations, which makes their impact on fitness difficult to predict. Epistasis can take place among genes or within genes (*3*), including between mutations that alter different features of a gene such as its regulation (or expression level, used here as equivalent to promoter activity) and others that alter protein activity or stability (coding). Unfortunately, regulatory and coding mutations are rarely studied simultaneously, apart from a few cases (*4*, *5*), such that we know little about how expression level modifies the fitness effects of coding mutations, and what the underlying mechanisms of such regulatory-by-coding epistasis would be. For instance, the relative effects of coding mutations could be amplified at low expression level. Conversely, mutations that are deleterious at low expression levels could become neutral at higher expression levels, or vice-versa. Thus, interactions between coding mutations and changes in gene expression could represent a vast unexplored constraint on protein evolution.

### The fitness landscape of a protein complex depends on its expression level

We investigated the extent to which gene expression level impacts on the fitness landscape of a protein by mimicking evolutionary changes by mutations affecting promoter activity using an inducible system. The bacterial dihydrofolate reductase DfrB1 is a homotetrameric enzyme with a single active site (*6*, *7*) that is inherently resistant to the antibiotic trimethoprim (TMP) (Fig. 1A-1B) (DfrB1 Ki ∼ 0.15 mM; *Escherichia coli* endogenous dihydrofolate reductase (ecDHFR) Ki ∼ 20 pM) (*8*, *9*). The fitness of *E. coli* can be made dependent on DfrB1 activity through the inhibition of ecDHFR using TMP. We used a plasmid whereby expression of *dfrB1* is regulated by an arabinose-dependent promoter (Fig. 1B). By inducing expression in 0 to 0.4 % arabinose, we found that WT DfrB1 expression at 0.2 % arabinose completely complements the inhibition of ecDHFR. This induction level was defined as the optimal expression level for WT DfrB1 (Fig. 1C, Fig. S1). Given the degree of complementation at each induction level, we refer to these expression levels as overexpressed (0.4 % arabinose), optimal (0.2 % arabinose), near-optimal (0.05 % arabinose), suboptimal (0.025 % arabinose) and weak (0.01 % arabinose) for the WT enzyme. We measured the corresponding expression levels by fusing DfrB1 to GFP (Fig. 1D). We observed a unimodal expression level within this range of induction. Our experiments show that DfrB1 has a typical enzymatic fitness function with increasing fitness with expression and then saturation (*10*).

**Fig. 1.**
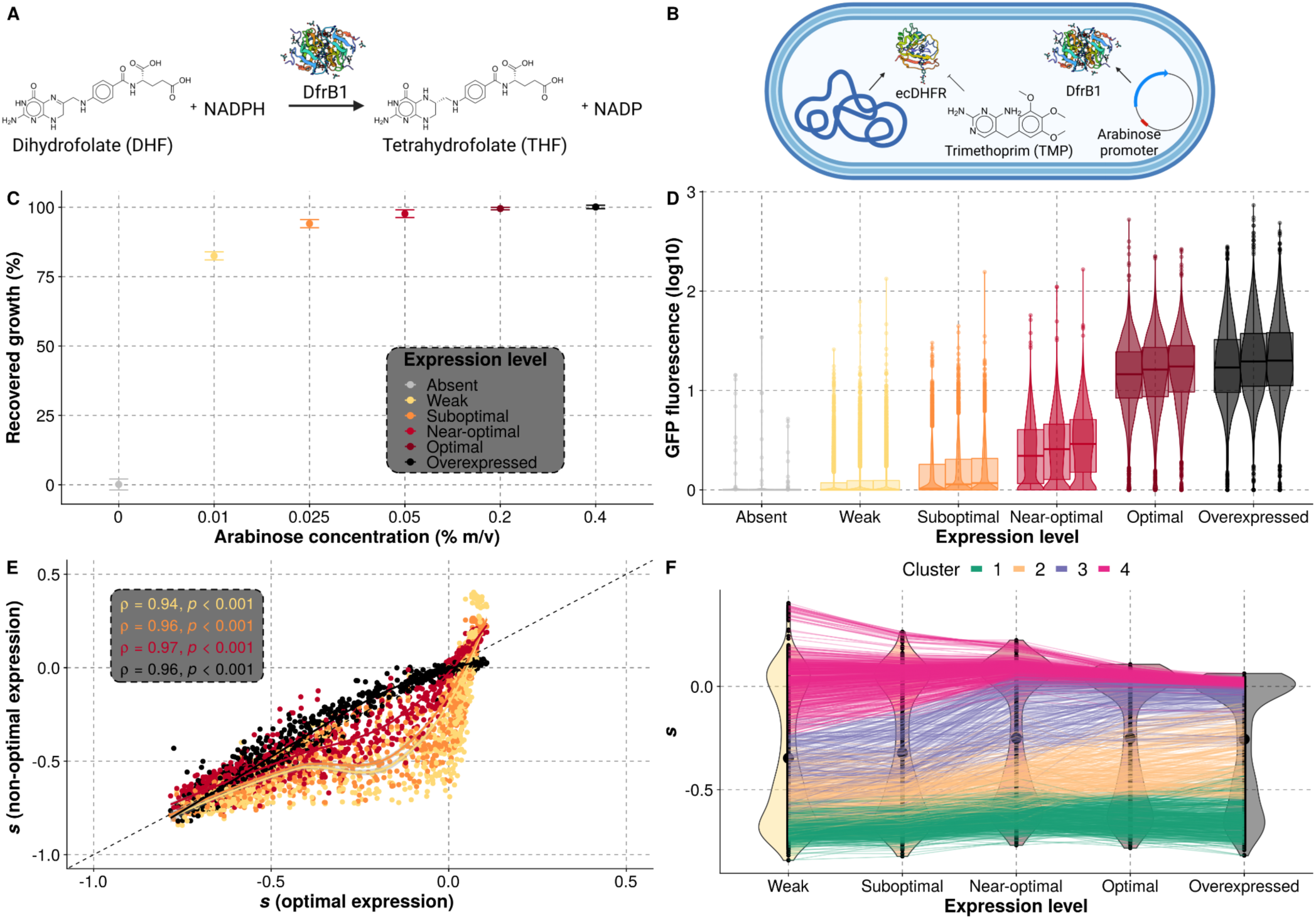
Selection coefficients (*s*) of amino acid changes depend on expression level. (**A**) DfrB1 converts dihydrofolate to tetrahydrofolate, an essential metabolite. (B) *E. coli* dihydrofolate reductase (ecDHFR) is inhibited by trimethoprim (TMP). DfrB1 is insensitive to TMP, and its transcription is modulated by an arabinose promoter. (C) Fitness-function of DfrB1 expression level. By inhibiting the endogenous ecDHFR with TMP (10 µg/mL), growth depends on DfrB1. Fitness is measured as a relative difference in growth (area under the curve) of the strain in medium with TMP relative to the same medium without TMP for a given arabinose concentration . Dots indicate the mean of six replicates and error bars indicate 2.5 standard errors of the mean. (D) DfrB1-GFP expression as a function of arabinose concentration. Expression is shown relative to an *E. coli* strain not expressing GFP. (E) Selection coefficient (*s*) of individual amino acid mutants at non-optimal levels of expression compared to the optimal level. Spearman correlation coefficients are shown. Lines indicate LOESS regressions. (F) *s* for individual mutants across expression levels. Colored lines indicate mutants from the different clusters (k-means, k = 4).

We generated a library of all possible single amino acid changes to the sequence of DfrB1 covering amino acid positions 2 to 78 (Fig. S2). Individual mutants were combined in a pool and transferred to medium with TMP at either of the five arabinose concentrations defined above (Fig. S3). The frequency of the mutants in the initial pool before selection and after 10 generations where DfrB1 function was required for growth was measured by deep sequencing (Table S1, Table S2). Redundant codons and replicates were aggregated to estimate a selection coefficient for each protein variant at each induction level (Fig. 1E-1F, Table S3). Biological replicates were highly correlated across experiments (Fig. S4). We confirmed the deleterious effects of previously reported mutants (Fig. S5, Fig. S6), and we validated a number of isolated mutants (n = 9) through growth curves (Spearman ⍴ = 0.87, p < 0.001) (Fig. S7).

The selection coefficients (*s*) of individual mutants are broadly correlated between expression levels, revealing that most amino acid substitutions have effects that are relatively similar across expression levels, but the correlation decreases as expression level deviates from optimal value (Fig. 1E). Overall, the average fitness effect of amino acid substitutions at a given position decreases as expression approaches the optimal level (Fig. S8A). A large fraction of substitutions (528 out of 1463 (36.1 %)) showed statistically significant effects of expression on fitness (magnitude > 0.1, ANOVA, FDR adjusted p < 0.05, Fig. S8B, Table S4). Clustering identified four typical patterns (k = 4 clusters) of selection coefficients along the expression gradients as the best compromise between parsimony and interpretability (Fig. 1F, Table S5, Fig. S9A,B). As one could expect, some substitutions do not have major effects on fitness (cluster 4) while others are deleterious at all expression levels (cluster 1). Interestingly, within cluster 4, some amino acid changes are advantageous at the lowest expression level (see below). Many variants appear to deviate from these two patterns (clusters 1 and 4), particularly between optimal and weak expression levels (Spearman ⍴ = 0.94) (Fig. 1E, Fig. S10). Mutants in cluster 3 are deleterious at weak and suboptimal expression, but they become neutral at the optimal expression level for the WT. Mutants from cluster 2 are even more deleterious at low expression levels but, even if they remain deleterious, they improve in fitness as expression increases, even beyond optimal expression for the WT. This gain of fitness of many mutants at expression level higher than optimal is also visible in the relationship between the selection coefficients at non-optimal versus optimal expression levels in Fig. 1E. The relationship is convex for expression levels lower than optimal for WT and slightly concave when overexpressed. Our results show widespread epistasis between expression level and coding mutations, with different expression levels favoring distinct mutants.

We also examined whether mutants changed fitness ranking among expression levels. An ANOVA on the ranks revealed significant effects of the interaction between genotype and expression level (F = 11.57, p < 2.2e-16), although these effects are much smaller than those associated to genotypes alone (F = 644.23, p < 2.2e-16) (Table S6). To validate this change in ranking, we grew more mutants (n = 8) individually. As previously, mutants with higher fitness at optimal expression for WT were clearly reproduced. Mutants that were less deleterious at lower expression were not as consistent. This means that it is more challenging to measure precise selection coefficients for highly deleterious amino acid substitutions. This is caused by the fact that these mutations are strongly underrepresented at the final timepoints in pooled competition experiments. However, the differences in fitness between expression levels in the individual versus bulk competition assays were strongly correlated (Spearman ⍴ = 0.72, p = 0.037) (Fig. S11).

### The protein structure determines which amino acid changes have effects that depend on expression levels

We mapped selection coefficients of all amino acid substitutions to individual positions (Fig. 2). Changes at conserved positions tend to be more deleterious than in non-conserved regions when comparing to diversity across homologs (Entropy, Fig. 2 - top) or using predictions from patterns of molecular evolution (*11*) (Fig. S12). Selection coefficients therefore reflect the long term evolution of this enzyme. The fitness effects of amino acid changes are heterogenous along the length of the sequence and depend on the structural property of each segment. For instance, substitutions in the poorly conserved and disordered region (positions 2-21) (*7*, *12*) tend to be more neutral, which is consistent with the observation that the sequence essential for DfrB1 function begins between residues 16-26 (*13*). Importantly, mutants from specific protein sites and with different destabilizing effects tend to be enriched in different clusters (Fig. S9C), suggesting that expression-dependent fitness effects can be explained by the protein structure (see below). Comparison of the landscapes at the five expression levels confirms a dampening of color gradient from top to bottom, showing that extreme fitness effects become rarer as expression level approaches the WT optimum. This means that at expression levels near the optimum for the WT enzyme, many amino acid substitutions tend to have positive and negative effects on fitness that are buffered. A change of expression levels therefore changes which mutations become visible to selection.

**Fig. 2.**
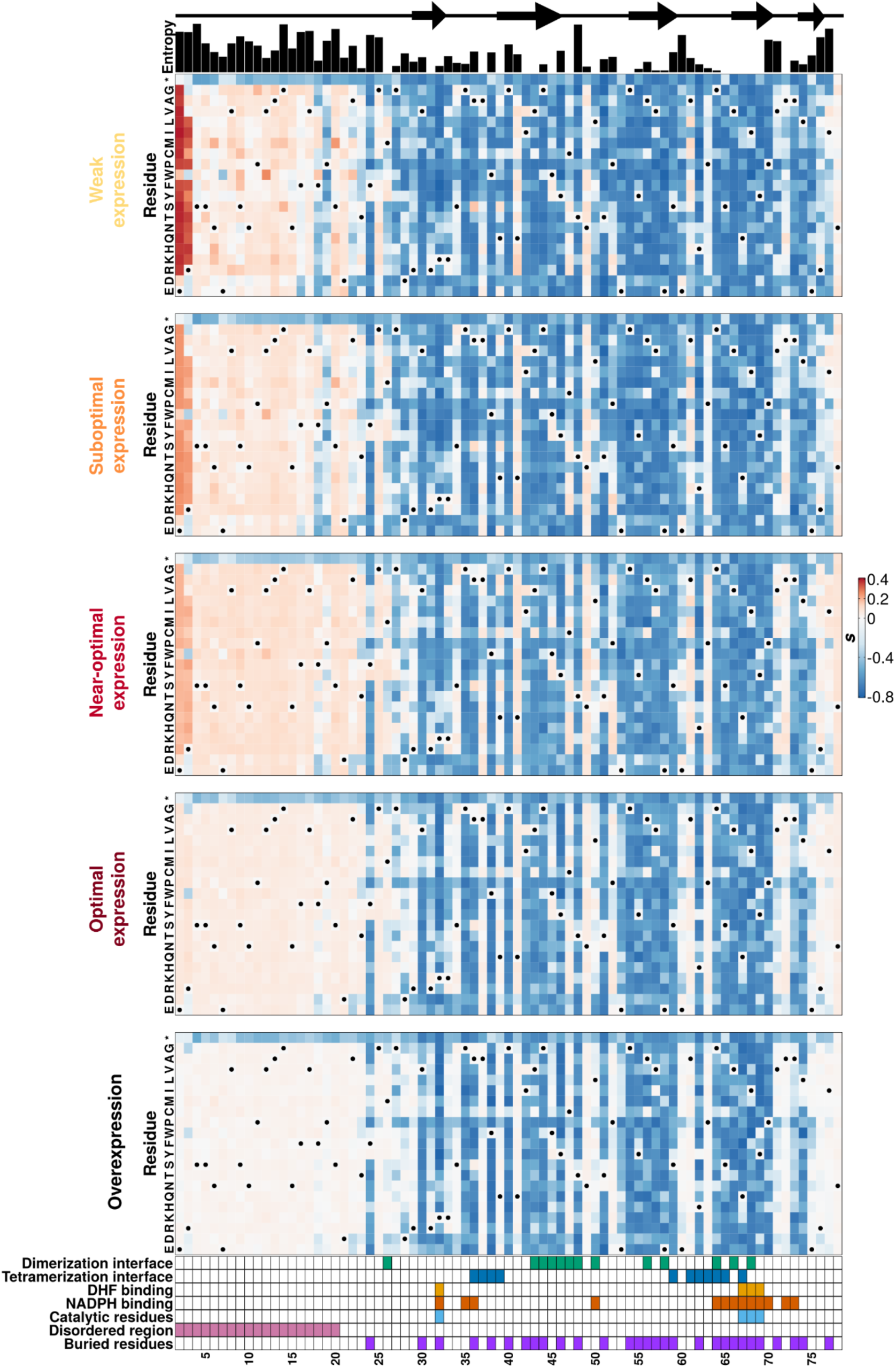
The DfrB1 fitness landscape changes with expression level in a way that depends on structural features. Selection coefficient (*s*) of individual amino acid mutants at five expression levels. Top annotations indicate the protein’s local secondary structure and sequence diversity among homologs (Shannon entropy). The bottom annotation highlights residues that participate in the interfaces of the DfrB1 homotetramer, that come into contact with either the substrate (DHF) or the cofactor (NADPH), and the key catalytic residues. Interfaces are labeled as dimerization and tetramerization interfaces (*14*). Residues marked with a dot in the heatmaps are wild-type residues (and their synonymous mutants).

### Masking of deleterious mutations is encoded in the protein structure

To capture the effect of expression level on selection coefficients, we calculated **Δ**s as the difference between the selection coefficients of an amino acid substitution at one of the non-optimal expression levels and at the optimal expression for the WT (**Δ***s_non-opt_ = s_non-opt_ -s_opt_*). A negative value indicates a more advantageous sequence at optimal expression. A positive value shows a more deleterious effect at optimal expression. A heatmap similar to Fig. 2 but with the **Δ***s_weak_* scores (Fig. 3A) shows that: 1) a majority of positions with deleterious mutations are deleterious no matter the level of expression (**Δ***s_weak_* around 0), 2) the deleterious effect of some mutations is dampened at optimal expression (**Δ***s_weak_* < 0), and 3) some mutations are beneficial specifically at low expression (**Δ***s_weak_* > 0). The dampening of mutational effects by gene expression is dose-dependent since the magnitude of **Δ***s* becomes smaller as expression level approaches the optimum (Fig. S13).

**Fig. 3.**
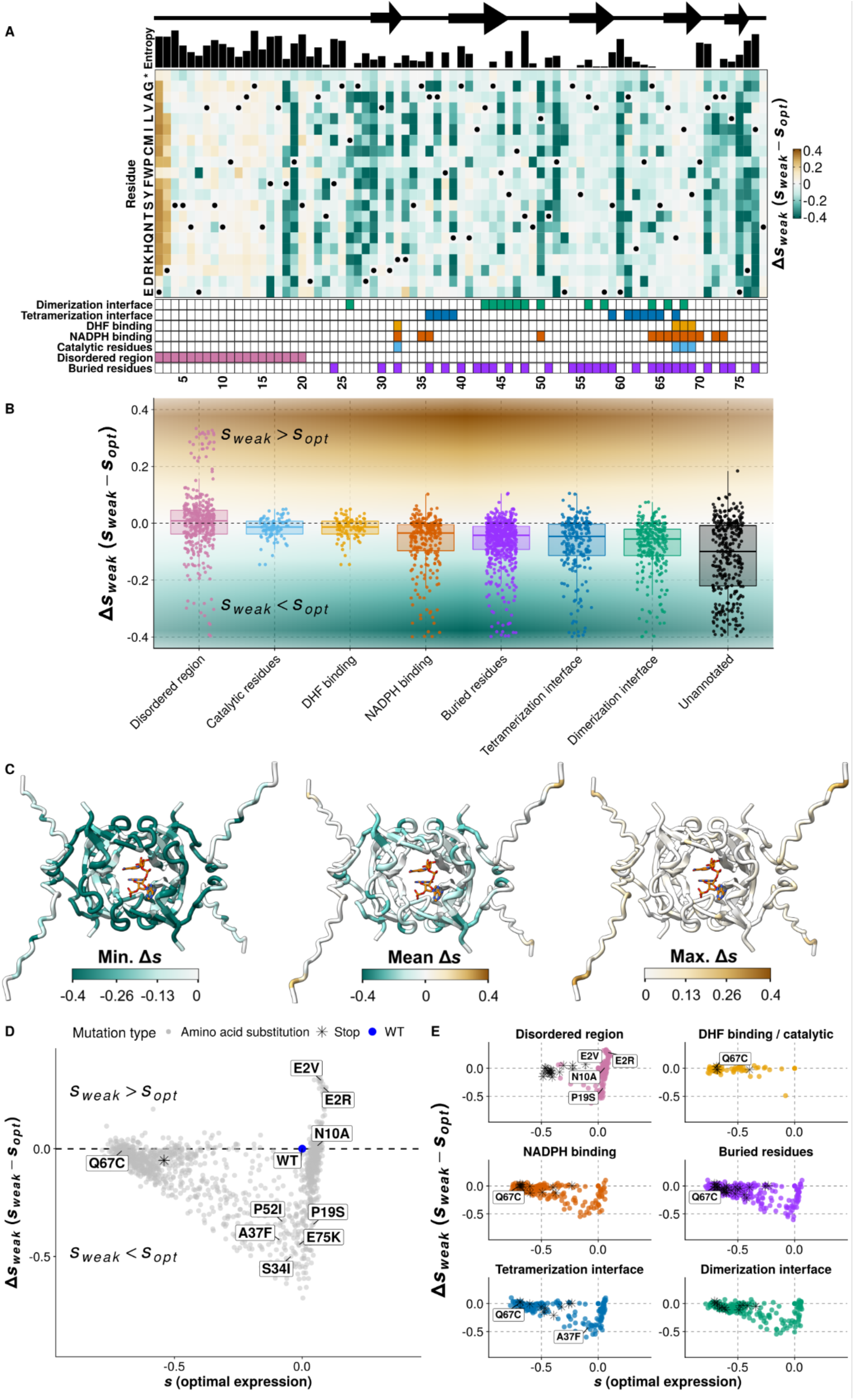
The deleterious effects of mutations can be buffered by WT optimal expression when they are slightly destabilizing but not when they affect catalytic activity. (**A**) **Δ***s* for weak versus optimal expression levels. A negative **Δ***s_weak_* value indicates mutations that are more advantageous at optimal expression for the WT. A positive **Δ***s_weak_* value indicates mutations that are more advantageous at weak expression. Annotations at the top (Shannon entropy) and the bottom (sites of interest in the protein) are the same as in Fig. 2. (B) Distributions of **Δ***s_weak_* vary with features of DfrB1 that correspond to structural properties. (C) Minimum, mean and maximum **Δ***s_weak_* values mapped on the DfrB1 structure. (D) Overall landscape of selection coefficients at optimal expression for the WT (*s_opt_*, x-axis) and **Δ***s_weak_* (y-axis). Mutants close to the dashed horizontal line have fitness effects that are independent from expression levels. Mutants from growth curve validations (Fig. S7,S8) are identified for reference, including the catalytic inefficient Q67C and the advantageous amino acid changes at weak expression E2R and E2V. (E) Functional sites of the protein occupy distinct regions of the selection landscape. Stop codons are labeled as black asterisks.

The disordered N-terminal region is distinct from the rest of the protein as it contains beneficial amino acid changes whose effects disappear at optimal expression (**Δ***s_weak_* > 0), particularly at positions 2 and 3 (Fig. S13). We assayed enzyme activity in cellular extracts for two mutants at position 2 and confirmed an increase in enzymatic activity for mutants E2R and E2V relative to the WT (Fig. S14A). We reasoned that the observed increase in activity could be due to higher protein abundance instead of an increase in catalytic efficiency given that position 2 is in the disordered region, far from the active site. Indeed, truncating DfrB1 at position 16 still produces an active enzyme, whereas truncating at position 26 destabilizes the protein by 2.5 kcal/mol and results in no measurable activity (*13*). Amino acid substitutions and nucleotide content in the first few codons of a protein are known to increase expression in *E. coli* (*15*), which makes this region a target for beneficial mutations when transcription levels are low. To test the effect of amino acid changes on protein expression, we generated GFP fusions with the entire sequence of one of such mutants (E2R), the WT disordered region (amino acids 1-25), and the disordered region with the E2R mutant. In both cases, we observed a higher protein abundance, even when only fusing the disordered fragment to GFP, reaching more than a 10-fold expression change relative to the WT sequence (Fig. S14B). Such a beneficial mutation at low expression therefore acts by increasing protein abundance, most likely by increasing translation rate.

Whereas the disordered region is particularly rich in positive **Δ***s_weak_* (Fig. 3B), mutation of the catalytic residues cannot be masked at optimal expression (**Δ***s_weak_* around 0). For instance, the Q67 mutant has a catalytic efficiency reduced by 100-fold with respect to the WT DfrB1 (*16*) and has a **Δ***s_weak_* near 0 (**Δ***s_weak_* = -0.008). On the other hand, interaction interfaces are particularly rich with mutations that are masked by optimal expression. Because **Δ***s_weak_* also has a zero value at sites that are completely insensitive to mutation, to better differentiate the effects, we mapped **Δ***s_weak_* values in a two dimensional space where they are plotted against s at optimal expression (Fig. 3D). This representation shows the striking difference between interaction interfaces and catalytic residues, where the effects of mutations are modulated by expression in one case and are expression-independent in the other (Fig. 3E). Mutations that destabilize the protein complex, such as those in buried sites (Fig. 3B-C), could be susceptible to epistasis between expression level and coding mutations, since an increase in the expression level could compensate for lower stability, following the law of mass action. Even the sites in the disordered region with negative **Δ***s_weak_* (F18 and P19) (Fig. 2, Fig. 3, Fig. S15) could be explained by interaction interfaces. AlphaFold2 (*17*, *18*) predicts residues F18 and P19 to be in contact with W45 (Fig. S15A-B), which is known to interact with the disordered N-terminal region (*19*). These analyses show the sharp contrast between mutants that depend on their structural and enzymatic roles: those with **Δ***s_weak_* near 0 being often involved in catalysis, those with positive **Δ***s_weak_* potentially increasing translation rate at the start of the coding sequence and those with negative **Δ***s_weak_* being often within protein interaction interfaces.

### Slightly destabilizing mutants are masked at optimal expression

To confirm the contribution of protein destabilization to the observed **Δ***s_weak_*, we computationally estimated the effects of amino acid changes on the protein and complex stability. Overall, we find a weak significant correlation between **ΔΔ***G* of subunit stability and **Δ***s_weak_* (Spearman ⍴ = 0.10, p-value = 4.3e-4). By mapping destabilizing effects in the *s_opt_* versus **Δ***s_weak_* landscape, we see that highly destabilizing mutations tend to cluster with poor catalytic activity (for instance Q67C) and stop-codon mutants (Fig. 4A). Similarly, for effects on the stability of individual subunits and protein interactions, higher expression levels tend to improve fitness at all levels of destabilization but to a very limited extent for **ΔΔ***G* greater than 2 kcal / mol, which largely behave like stop-codon mutants (Fig. 4B). This observation also helps explain the differences in the distribution of **Δ***s_weak_* between buried and unannotated sites (Fig. 3B,E) because mutations in buried sites tend to be more destabilizing than mutations in exposed sites. To confirm the predominant role of protein destabilization, we trained a random forest regressor (*20*) (see methods) on **Δ***s_weak_* using the predicted biophysical effects of amino acid changes, structural features, changes in amino acid properties based on the differences in 57 indexes from ProtScale (*21*), and the propensity of amino acids to be found in protein-protein interaction interfaces (*22*) (Table S7). While the global performance was modest (*R*^2^ (test set) = 0.32, *R*^2^ (5-fold cross-validation) = 0.38 ± 0.034 (standard error of the mean)), it identified relative solvent accessibility, changes in hydrophobicity and effects on subunit stability as the top features contributing to the predictions (Fig. S16). As a result, we conclude that the relation between destabilizing effects and expression level is nonlinear, with expression around the WT optimal expression only masking slightly destabilizing mutants (**ΔΔ***G* < 2 kcal / mol).

**Fig. 4.**
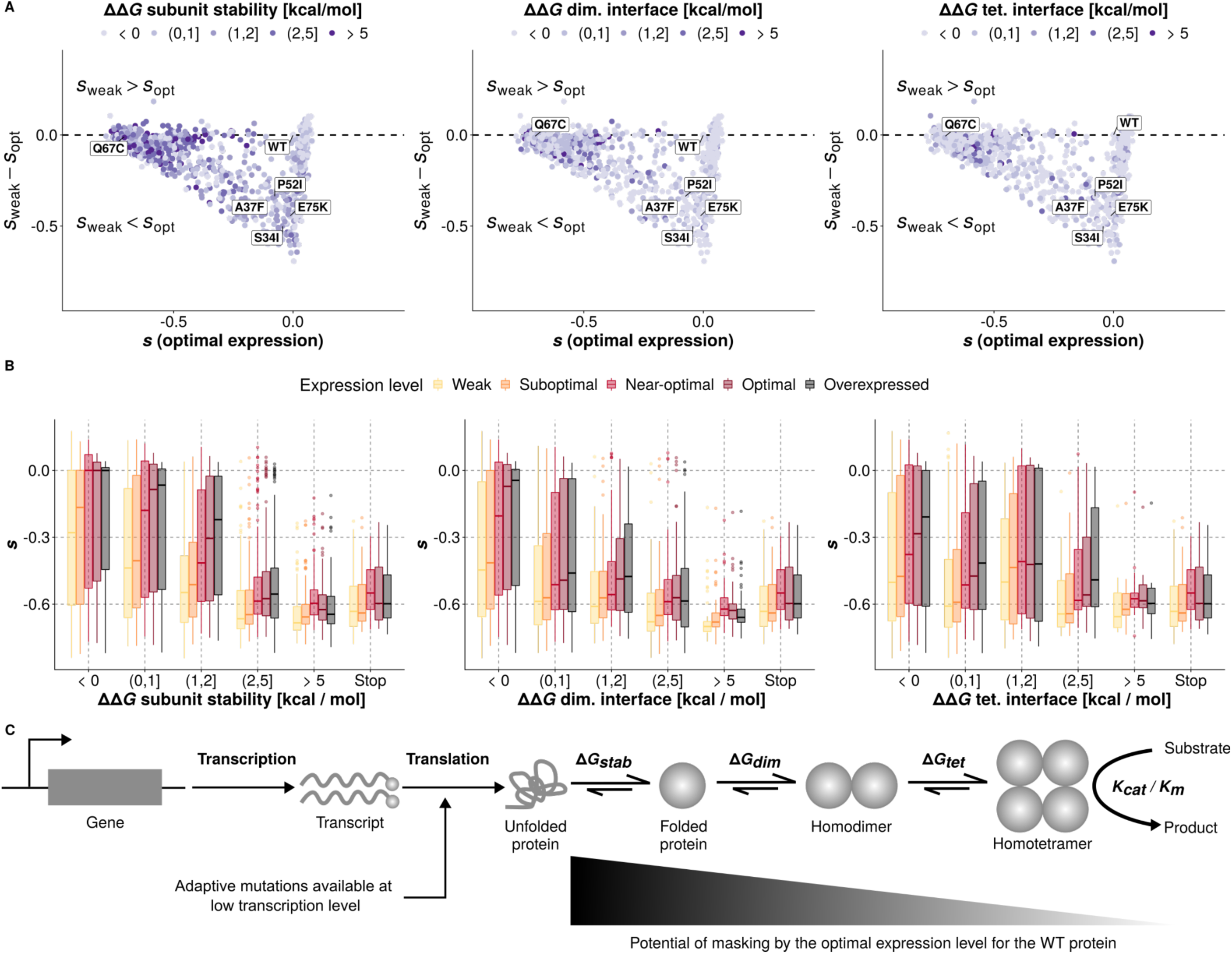
Optimal expression masks slightly destabilizing mutations. (**A**) Landscape of selection coefficients at optimal expression for the WT (x-axis) and the expression-dependent change in fitness effects (y-axis) labeled with respect to biophysical effects of mutations on subunit stability, the dimerization interface, and the tetramerization interface. Mutants at positions 2-20 were excluded from this analysis because their coordinates are not present in the PDB structure (2RK1). (B) Distributions of measured selection coefficients for mutants with different effects on protein stability or binding affinity at the interfaces. Stop codons from the functional protein core (residues 30 - 70) are shown for comparison. (C) Functional protein complex assembly is disturbed by mutations that can be masked by expression level. Mutations with small to moderate effects on protein stability and interface binding affinity can be masked by higher expression level, unlike mutations disrupting enzymatic active sites.

### Protein destabilization in itself does not reduce fitness

Destabilizing mutations could affect fitness in two ways. First, by reducing the amount of homotetrameric complex formed, which decreases enzyme activity. A second mechanism is the potential for misfolded proteins to create toxic aggregates and detrimental interactions with other proteins, which could be amplified by a higher expression level (*23*). Under this scenario, highly destabilizing mutations would critically reduce protein complex concentration at low expression levels and any gain resulting from higher expression level would be counterbalanced by the toxic effect of misfolding. Such amino acid substitutions would remain deleterious at all expression levels, as is shown in Fig. 1F for mutants in cluster 1 that are enriched for highly destabilized mutants.

This last hypothesis predicts that strongly destabilizing mutants would have a negative impact on fitness even when removing selection for DfrB1 activity. We first tested if expression of DfrB1 was toxic by measuring growth rate without TMP of the WT enzyme and of a construct lacking the start codon so no protein or much less protein is produced. We do not detect negative fitness effects in these assays (Fig. S17) which suggests that the WT protein itself is not toxic. However, to examine whether some mutants could be toxic, we repeated the bulk competition experiment of the mutant library, inducing expression in 0 to 0.4 % arabinose, this time removing selection for DfrB1 activity by omitting TMP. We observe lower signal-to-noise ratio, with a lower correlation between biological replicates (Fig. S18) than when TMP selected for DfrB1 activity. The distributions of selection coefficients are tightly clustered around 0, confirming the lack of detectable fitness effects of most mutations when selection is removed (Fig. S19, Fig. S20).

In spite of the very weak signals in these conditions, **Δ***s* values measured without TMP are weakly negatively correlated with the **Δ***s* values measured with TMP, particularly when DfrB1 is overexpressed (weak expression: Spearman ⍴ = -0.06, p = 0.014; suboptimal expression: ⍴ = -0.04, p =0.062; near-optimal expression: ⍴ = -0.006, p = 0.81; optimal expression: ⍴ = -0.07, p = 0.004; overexpression: Spearman ⍴ = -0.16, p = 8.79e-11). This would suggest that the most performing mutants in the presence of TMP have reduced fitness when expressed in the absence of a need for this enzyme. Consistent with this, while fitness effects in medium with TMP are lower for amino acid substitutions predicted to be deleterious based on molecular evolution of homologs, the correlation is inverted without TMP (Fig. S21). Fitness reduction in the absence of selection is therefore particularly noticeable for functional variants, which suggest that destabilization-based toxicity plays no or only a minor role. To further examine this, we analyzed the relation between **ΔΔ***G* and *s* (Fig. S22). We fitted generalized linear models (GLM) for the selection coefficients (*s*) observed with and without TMP using the destabilizing effects of mutations and their interactions with expression level (Table S8). The coefficients for destabilizing effects are all negative and statistically significant in the model for the data with TMP. However, only mutational effects on protein stability had a statistically significant effect in the GLM for the data without TMP and this coefficient was slightly positive (1.69e-4). Thus, we conclude that any deleterious effects caused by misfolding of DfrB1 in the absence of selection are minimal and that DfrB1 activity would instead reduce fitness when overexpressed due to metabolic effects. This is consistent with reports that an additional copy of a dihydrofolate reductase alters cell metabolism (*24*) and the fact that the only sites for which we found a deleterious effect without TMP at overexpression level are mutations in the second and third codons (Fig. S23) that lead to increases in protein abundance and total enzyme activity (Fig. S14). This leads to the prediction that the E2R mutant would have a lower optimal induction level than the WT. Indeed, we observed that the E2R mutant leads to a higher percentage of growth recovery than the WT at low induction levels but decays slightly at the WT optimal expression level and overexpression, revealing a metabolic cost of high DfrB1 abundance (Fig. S24).

## Conclusion

By measuring how expression level (used as being equivalent to promoter activity throughout) impacts the fitness effects of amino acid substitutions, we find that many slightly deleterious mutations tend to become less deleterious as expression approaches the optimal expression level of the wild-type protein. One key factor to explain this is the impact of mutations on protein stability rather than on the intrinsic enzyme activity. Coding mutations that increase protein abundance are beneficial at low expression level but become neutral as expression gets optimal for the WT sequence or even costly when optimal expression is overshot. Changes in gene regulation, for instance through alteration of promoter activity by mutation, can therefore dictate which mutations will be beneficial or deleterious for a coding sequence, and vice-versa (Fig. 4C). Over the long term, promoter activity and protein sequences will thus coevolve with each other. This co-evolution will be disrupted when major changes occur in the regulation of the expression level of a gene, such as in the event of gene duplication, genomic rearrangements, or horizontal gene transfer. Such events, if they lead to expression levels that are far from optimal, could result in bursts of adaptive protein evolution by changing which mutants become targets for selection and this, without improving the protein functions.

Our results may seem counterintuitive given the observation that highly expressed proteins evolve slowly (*25*). If high expression buffers the effect of mutations, one would wrongly expect highly expressed proteins to evolve faster. However, since highly expressed proteins are costly to produce (*26*), their expression level may actually be below the optimal level that would be observed in the absence of metabolic cost. As a result, their fitness landscape would be more similar to the fitness landscapes we observed at weak and suboptimal expression, where purifying selection against destabilizing mutations would be more intense, thereby preserving protein sequence.

## Acknowledgements

We thank C. Lemay-St.Denis, D. I. Ascencio, R. Durand, M. Dion, F. Mattenberger and the Landrylab for feedback and help on the project and M. Couture for plasmids. Figures 1A-B and S3 were generated using Biorender.

## Funding

Canadian Institutes of Health Research (CIHR) Foundation grant to 387697 (CRL)

NSERC discovery grant RGPIN-N-2018-04686 (JNP)

Canada Research Chair in Cellular Systems and Synthetic Biology (CRL)

Canada Research Chair in Engineering of Applied Proteins (JNP)

FRQNT Merit Scholarship Program for Foreign Studies (PBEEE) (AFC)

Joint funding from MEES and AMEXCID (AFC)

Vanier graduate scholarship (PCD)

## Contributions

Conceptualization: AFC, IGA, CRL

Data curation: AFC, IGA, KL

Investigation: IGA, AKD, KL and AFC

Formal analysis: AFC, IGA and CRL

Software: AFC, PCD, and PK

Writing – original draft: AFC, CRL, and IGA

Writing – review & editing: AFC, CRL, IGA, PCD, AKD, KL, JNP

Supervision: JNP and CRL

Funding acquisition: JNP and CRL

Methodology: AFC, IGA, CRL, KL, JNP

Project administration: CRL

Resources: CRL

Validation: AFC, IGA

Visualization: AFC

**Competing interests:** Authors declare that they have no competing interests.

**Data and materials availability:** Raw sequencing data are available at SRA BioProject PRJNA842350 (accession numbers SRR19419448 and SRR19419449). Code for analysis is available on Github at: https://github.com/Landrylab/DfrB1_DMS_2022

All data, code, and materials used in the analysis are available in the supplementary files, and dataset on through SRA. All the material is available upon request or from Addgene for the pBAD-sfGFP plasmid.

## Supplementary Materials

## Materials and Methods

All strains, reagents, compounds and softwares used in this study are listed and referenced in Table S9.

### 1.- Strains, media and plasmids

MC1061 is the *E. coli* strain used for all cloning and mutagenesis steps, whereas *E. coli* BL21 (DE3) is the one used for the experiments conducted in this study. Transformations with plasmids were done according to standard procedures with in-house chemically competent cells (*27*). Transformed bacteria were grown and selected on 2x YT + glucose medium (1.0 % yeast extract, 1.6 % tryptone, 0.2 % glucose, 0.5 % NaCl and 2 % agar; (*28*)) with ampicillin (AMP; 100 µg/mL). For all experiments conducted in liquid medium (with the exception of the growth to measure DfrB1 activity; see below), bacteria were grown in LB (Luria-Bertani) medium (0.5 % yeast extract, 1.0 % tryptone, 1.0 % NaCl; (*29*)) with AMP and with or without L(+)-arabinose (0.001 to 0.4 %) and trimethoprim (TMP; 10 µg/mL in DMSO). To measure DfrB1 activity, bacteria were grown in TB (Terrific broth) medium (2.4 % yeast extract, 1.2 % tryptone, 0.4 % glycerol and 89 nM potassium phosphate).

The *dfrB1* gene was expressed in bacteria from a pBAD vector which allows arabinose-controlled induction (*30*). pBAD-*dfrB1* was constructed as follows. The pBAD vector was amplified from pBAD-*chuA* plasmid (*31*) with CLOP198-E9 and CLOP198-F9 (PCR program: 5  min at 95  °C, 35 cycles: 20  s at 98  °C, 15  s at 61  °C, 2 min 30 at 72  °C, and a final extension of 3  min at 72  °C). *dfrB1* was amplified from pDNM plasmid with CLOP198-A10 and CLOP198-B10 (PCR program: 5  min at 95  °C, 5 cycles: 20  s at 98  °C, 15  s at 62  °C, 15 s at 72  °C, 30 cycles: 20  s at 98  °C, 15  s at 72  °C, 15 s at 72  °C, and a final extension of 3  min at 72  °C). Both PCR products were incubated for 1 h at 37 °C with 20 U of DpnI enzyme to remove parental DNA and subsequently purified on magnetic beads (Axygen AxyPrep Mag PCR Clean-up Kit). The vector (pBAD) and insert (*dfrB1*), with their overlapping regions at each extremity, were then assembled by Gibson DNA assembly (*32*), producing pBAD-*dfrB1* plasmid.

To measure by cytometry the effect of mutating the second residue of DfrB1 on the expression level, we fused GFP in 3’ of the *dfr*B1 gene sequence and of its mutant. The plasmid was amplified from pBAD-*dfrB1* (constructed above) or from pBAD-*dfrB1*(E2R) (isolated from the DMS plasmid collection -see below) with CLOP273-A3/B3 (PCR program: 5  min at 95  °C, 35 cycles: 20  s at 98  °C, 15  s at 61  °C, 2 min 30 at 72  °C, and a final extension of 3  min at 72  °C). GFP was amplified from pBAD-sfGFP(*33*) with CLOP273-D3/F3 (PCR program: 5  min at 95  °C, 5 cycles: 20  s at 98  °C, 15  s at 62  °C, 20 s at 72  °C, 30 cycles: 20  s at 98  °C, 15  s at 72  °C, 20 s at 72  °C, and a final extension of 3 min at 72  °C). The PCR products corresponding to the plasmids and insert were incubated for 1 h 30 at 37 °C with 10 U of DpnI enzyme to remove parental DNA. Subsequently, the plasmids (pBAD-*dfrB1* and pBAD-*dfrB1*(E2R)) and insert (*sfGFP*) were purified on magnetic beads (Axygen AxyPrep Mag PCR Clean-up Kit) and assembled by Gibson DNA assembly (*32*), producing pBAD-*dfrB1*-*sfGFP* and pBAD-*dfrB1*(E2R)-*sfGFP*.

We also fused the GFP to a fragment corresponding only to DfrB1 first 25 residues. Vectors and the *sfGFP* insert were amplified from the same plasmids as above but using respectively CLOP273-A3/C3 or CLOP273-E3/F3 primers. After DpnI digestion and magnetic beads purification, Gibson DNA assembly produced pBAD-*dfrB1*[1-25]-*sfGFP* and pBAD-*dfrB1*[1-25](E2R)-*sfGFP* plasmids.

To insert specific mutations in the *dfrB1* sequence, we performed site-directed mutagenesis based on the QuickChange™ Site-Directed Mutagenesis System (Stratagene, La Jolla, CA). Briefly, we amplified the pBAD-*dfrB1* plasmid using pairs of primers containing the desired mutation at the center (see Table S10 for the primers specific to each mutation) (PCR program: 2  min at 95  °C, 22 cycles: 20  s at 98  °C, 15  s at 68  °C, 3 min at 72  °C, and a final extension of 5  min at 72  °C). The PCR products were then incubated for 1 h 30 at 37 °C with 6 U of DpnI enzyme to remove parental DNA and mutated plasmids were retrieved directly by transformation in bacteria. We used this method to generate the following mutants: pBAD-*dfrB1*(F24V), pBAD-*dfrB1*(K33E), pBAD-*dfrB1*(K33L), pBAD-*dfrB1*(K33M), pBAD-*dfrB1*(S59Y) and pBAD-*dfrB1*(I68M).

To generate a mutant in which the first methionine codon of DfrB1 is replaced by a stop codon (**Δ**MET mutant -pBAD-*dfrB1*(**Δ**MET)), we performed a PCR reaction using 10 ng of pBAD-*dfrB1* plasmid as a template (PCR program: 5  min at 95  °C, 25 cycles: 20  s at 98  °C, 15  s at 59  °C, 2 min 30 at 72  °C, and a final extension of 3  min at 72  °C) with the non-overlapping primers (CLOP265-A1 and CLOP259-H5) designed to incorporate a TGA stop codon on the forward primer to replace the AUG. The PCR product was then digested at 37 °C for 1 hour with 10 U DpnI. The digested PCR product was purified on magnetic beads, quantified and ∼50 ng were taken for phosphorylation using 5 U of T4 PNK in T4 ligase buffer for 30 min at 37 °C (final reaction volume of 5 µL). Subsequently, ligation was done in a reaction volume of 10 µL by adjusting the volume of T4 ligase buffer, adding 10 U of T4 DNA ligase and incubating for 1 hour at 22 °C. Finally, mutated plasmid was retrieved directly by transformation in *E. coli* BL21 bacteria.

For fitness validation, some of the plasmids that were used were directly isolated from the mutagenesis pools but confirmed by sequencing. Briefly, an aliquot of glycerol stock for the position of interest was diluted and plated on solid medium. Colony PCR was performed on 32 colonies to amplify the *dfrB1* gene using CLOP228-C1/D1 (PCR program: 5  min at 94  °C, 30 cycles: 30  s at 94  °C, 30  s at 54  °C, 35 s at 72  °C, and a final extension of 3  min at 72  °C). Two subsequent PCR rounds were used to add specific row-column and plate barcodes (*34*). These PCRs are described in detail in the library sequencing section below. All pooled samples were sent to the Genomic Analysis Platform (IBIS, Quebec, Canada) for paired-end 300  bp sequencing on a MiSeq (Illumina). After identification of individual mutations, plasmids for clones of interest were retrieved and purified from bacteria using the Presto Mini Plasmid Kit.

All PCR reactions mentioned above or described below were performed with oligonucleotides defined in (Table S10), using Kapa polymerase at the exception of the colony PCRs for which Taq DNA polymerase was used. The integrity of all assembled and mutagenized plasmids was confirmed by Sanger sequencing (Plateforme de séquençage et de génotypage des génomes, CRCHUL, Canada) using either CLOP194-G11 or CLOP196-B5 as a sequencing primer.

### 2.- Deep mutational scanning

A single-site mutation library of *dfrB1* was generated by a PCR-based saturation mutagenesis method (*35*). The mutagenesis was carried out in a codon-position-wise manner, directly on the pBAD-*dfrB1* plasmid with the oligonucleotides defined in (Table S10). Forward oligonucleotides which have degenerate nucleotides (NNN) so each codon of *dfrB1* is mutated were used in combination with a single reverse primer located in the plasmid outside the coding sequence. A first PCR to generate an amplicon containing the desired mutations was conducted following these steps: 5  min at 95  °C, 35 cycles: 20  s at 98  °C, 15  s at 60  °C, 30  s at 72  °C, and a final extension of 1  min at 72  °C. The resulting PCR product was then used as a mega primer to introduce the mutations in pBAD-*dfrB1* by amplifying the whole plasmid (PCR program: 5  min at 95  °C, 22 cycles: 20  s at 98  °C, 15  s at 68  °C, 5 min at 72  °C, and a final extension of 7  min at 72  °C). The long PCR product was digested for 90 min at 37  °C with 6 U of DpnI to remove parental DNA. The digestion products for individual positions were transformed into *E. coli* MC1061. More than a thousand colonies were retrieved from each transformation. After addition of glycerol, libraries for each position were stored separately. From the same pools of bacteria, plasmids were also extracted and purified. PCR amplification and Illumina MiSeq sequencing, performed on these plasmids allowed library quality control assessment (Fig. S2). In an initial round, mutagenesis of position 39 was not successful. Mutagenesis at this position was repeated separately and added to the final library after quality control.

### 3.- Bulk competition assay

First, *E. coli* BL21 was transformed with 75 ng of each individual position mutant pool (DNA miniprep from the DMS step above). Start and stop codon positions (positions 1 and 79) were omitted. All colonies were retrieved from each transformation plate in 5 mL of 2x YT liquid medium. OD_600_ was measured for each pool (final *OD*_600_ between 40-80 after being scraped and resuspended in medium). At this step, 15 % glycerol was added to the medium and an aliquot of each pool was stored individually for further use. In parallel, pools were equally mixed at an OD_600_ of 25 to generate a starting pool for the bulk competition assay. This master pool was stored for further use.

Two separate bulk competition assays were performed. Details concerning arabinose concentrations, the presence or absence of TMP and the number of replicates are indicated in Table S1. Briefly, the master pool was used to inoculate at *OD*_600_ = 0.01 a first preculture in LB + AMP medium. After an overnight incubation at 37 °C with agitation (250 rpm), cells were diluted 1:100 in fresh medium with the addition of different amounts of arabinose. Following an 18-hour incubation at 37 °C (250 rpm), cells were diluted again at *OD*_600_ = 0.025 in a final volume of 4 mL of fresh medium containing arabinose with the addition of TMP or DMSO (TMP solvent - no TMP control). Cultures were then incubated as above until *OD*_600_ reached 0.8 (5 generations). At this point, 125 µL was used to dilute back to *OD*_600_ = 0.025 in fresh medium and cells were grown until once again, *OD*_600_ reached 0.8 (another 5 generations). Finally, at two timepoints (18-hour preculture = Timepoint 0, after 10 generations = Timepoint 10), 3 mL of the culture was used to extract plasmid DNA (Fig. S3). These samples were treated as described below to prepare libraries for sequencing.

### 4.- Single mutant library sequencing

Libraries for sequencing were generated as described in Dionne et al. (*36*). Briefly, three PCR steps were done. The first one was performed directly on the small plasmid DNA preparations (4.5 ng of plasmid) corresponding to single-codon mutant libraries or to bulk competition assay samples (PCR program: 3  min at 98  °C, 20 cycles: 30  s at 98  °C, 15  s at 60  °C, 30  s at 72  °C, and a final extension of 1  min at 72  °C). The second PCR was performed to add Row and Column barcodes (*34*) for identification in a 96-well plate (PCR program: 3  min at 98  °C, 15 cycles: 30  s at 98  °C, 15  s at 60  °C, 30  s at 72  °C, and a final extension of 1  min at 72  °C). For this second PCR, the first PCR product was used as a template (2.25  µL of a 1/2500 dilution). Quantification on gel of the second PCR product using Image Lab (BioRad Laboratories) allowed us to mix the libraries so each has a roughly equal amount in the final library. Mixed PCRs were purified on magnetic beads and quantified using a NanoDrop (ThermoFisher). Finally, 0.0045 ng of the purified mixed PCRs were used as template for the third PCR which adds a plate barcode as well as Illumina adapters (PCR program: 3  min at 98  °C, 18 cycles: 30  s at 98  °C, 15  s at 61  °C, 35  s at 72  °C, and a final extension of 1  min at 72  °C). Each reaction for the third PCR was performed in quadruplicate and then combined. After purification on magnetic beads, libraries were quantified using a NanoDrop (ThermoFisher) and sent to the Genomic Analysis Platform (IBIS, Quebec, Canada) for paired-end 300  bp sequencing on a MiSeq (Illumina) or to the Plateforme de séquençage et de génotypage des génomes (CRCHUL, Quebec, Canada) for paired-end 250 bp sequencing on a NovaSeq (Illumina). All raw data are available at SRA BioProject PRJNA842350 (accession numbers SRR19419448 and SRR19419449).

### 5.- Bacterial growth curves

To measure fitness for individual WT or mutant DfrB1, growth was followed with serial *OD*_600_ measurements in a plate reader. From an overnight preculture grown in LB + AMP medium, cells were diluted 1:100 in fresh medium with the addition of 0 % to 0.4 % arabinose according to the experiment. Following an 18-hour incubation at 37 °C (250 rpm), cells were diluted again at *OD*_600_ = 0.01 in a final volume of 200 µL of fresh medium containing arabinose with the addition of TMP or DMSO. The 96-well plate was incubated at 37 °C in an Infinite M Nano plate reader (Tecan) for 20 hours. *OD*_600_ measurements were taken every 15 min. Plate was agitated at 200 rpm in between measurements. Growth curves were analyzed with the Growthcurver R package (*37*) to calculate the area under the curve as an estimate of fitness. The percentage of recovered growth upon arabinose induction was calculated as follows:

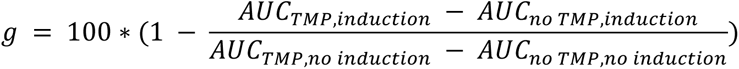

where g refers to the recovered growth percentage and *AUC* refers to the area under the curve in the conditions specified by the subscript (with or without TMP, with or without arabinose to induce expression).

### 6.- Expression level measurement by flow cytometry

To measure GFP level by cytometry, a first preculture in LB + AMP medium was grown with bacteria containing the plasmid of interest (pBAD-*sfGFP*, pBAD-*dfrB1*-*sfGFP*, pBAD-*dfrB1*(E2R)-*sfGFP*, pBAD-*dfrB1*[1-25]-*sfGFP* and pBAD-*dfrB1*[1-25](E2R)-*sfGFP*). As for the bulk competition assay, after an overnight incubation at 37 °C with agitation (250 rpm), cells were diluted 1:100 in fresh medium with the addition of different amounts of arabinose. Following an 18-hour incubation at 37 °C (250 rpm), cells were diluted again at *OD*_600_ = 0.025 in a final volume of 4 mL of fresh medium containing arabinose. Cultures were then incubated as above until *OD*_600_ reached 0.8 (5 generations). At the different timepoints (18-hour preculture = Timepoint 0, after 5 generations = Timepoint 5), small aliquots of cells were taken and diluted in sterile filtered water to an *OD*_600_ = 0.05 in 200 µL. GFP fluorescent measurements as well as forward scatter (FSC) and side scatter (SSC) data were collected from a Guava easyCyte HT cytometer (Luminex). From the cytometry data, *E. coli* cells were selected based on FSC and SSC. From the selected data points, GFP fluorescence signal was measured after excitation with a blue laser (wavelength = 488 nm) and detection with a green channel (525/30 nm).

### 7.- DfrB1 activity measurement

Enzymatic activity of DfrB1 and of different mutants was tested *in vitro*. From a 16-18 h 5 mL LB + AMP preculture (incubation at 37 °C with agitation (230 rpm), a 10 mL culture in TB + AMP was inoculated at *OD*_600_ = 0.1 and incubated for 3 h at 37 °C (OD_600_ = 0.7 to 1) with agitation. Induction of expression was initiated by addition of 1 % arabinose and incubation was continued at 22 °C for 16-18 h with agitation. After induction, cells were pelleted for 30 min at 3000 rpm (Eppendorf Centrifuge 5810 R) at 21 °C, resuspended in 300 µL of lysis buffer (0.1 M KH_2_PO_4_-K_2_HPO_4_ pH 8.0, 10 mM MgSO_4_, 1 mM DTT, 0.5 mg/mL lysozyme, 5 U DNase, 1.5 mM benzamidine, 0.25 mM PMSF) and incubated for 2 h at 30 °C with vigorous shaking. Lysates were cleared by a 30 min centrifugation at 3000 rpm (Eppendorf Centrifuge 5810 R) at 21 °C. In a 96-well plate, cleared lysates (20 µL) were combined with 50 µg/mL TMP, 100 µM DHF (synthesized as in (*38*)) and 100 µM NADPH in 50 mM KH_2_PO_4_-K_2_HPO_4_ pH 7.0 in a total volume of 100 µL. Measurement of absorbance at 340 nm was taken for 5 min in a Beckman Coulter DTX880 Multimode detector plate reader. Enzyme activity was calculated using the slope for the initial 20 % of substrate consumption.

### 8.- Data analyses

#### 8.1.- Inferring fitness scores from sequencing data

Quality control of the MiSeq and NovaSeq sequencing data was performed using FastQC version 0.11.4 (*39*). Trimmomatic version 0.39 (*40*) was used to select reads with a minimal length from the raw data (MINLEN parameter: 299 for MiSeq reads, 250 for NovaSeq reads) and trim them to a final length (CROP parameter: 270 for MiSeq reads, 225 for NovaSeq reads) . Selected reads were aligned using Bowtie version 1.3.0 (*41*) to the plate, row, and column barcodes to demultiplex sequences from each pool (arabinose concentration + timepoint). The remaining paired reads were merged with Pandaseq version 2.11 (*42*) and identical sequences were aggregated using vsearch version 2.15.1 (*43*). Finally, aggregated reads were aligned to the reference sequence of DfrB1 to identify the mutant codon in each read. Stop codon UAG was removed from all analysis because it has been shown to have a lower termination efficiency than the other stop codons in *E. coli (44, 45)*.

Raw read counts were normalized by the total number of reads in each pool (arabinose + timepoint). The resulting read proportions were used to calculate selection coefficients based on the end point and the starting point of each experiment using the following equation:

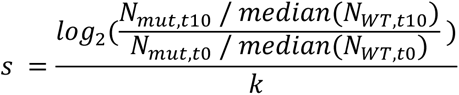

where s is the selection coefficient, N is the number of reads for the corresponding mutant at a specific timepoint, and k is the number of generations (k = 10).

#### 8.2.- Evolutionary analysis (GEMME, Evol, jackhmmer)

A set of seven DfrB1 homologs was extracted from data published by Toulouse et al. (*46*). These seven sequences were then submitted to the online jackhmmer tool (https://www.ebi.ac.uk/Tools/hmmer/search/jackhmmer) (*47*) using the Uniprot database as a target to look for additional homologous sequences. The resulting set of sequences were filtered with the following criteria, considering that the full-length DfrB1 sequence is 78 residues long:

- E-score < 1e-6

- Alignment length > 50

- Sequence length < 100

Applying these filters resulted in a final set of 82 high-confidence DfrB1 homologs. These sequences were aligned with MAFFT version 7.475 (*48*, *49*) using the iterative refinement method (*50*, *51*). The resulting alignment was then analyzed with the Evol library from the ProDy suite version 2.0 (*52*) to calculate Shannon entropy at each position in the alignment as a metric of evolutionary variation. In parallel, the same MAFFT alignment was analyzed with the GEMME model (*11*) to obtain predictions of the fitness landscape based on the variation in the alignment.

#### 8.3.- Machine learning analyses 8.3.1.- k-means clustering

We used k-means clustering to identify the typical patterns of expression-dependent changes in fitness effects (Fig. 1F). k-means clustering was run using the kmeans function from the R base stats package (*53*) by setting the seed and using values of k from 1 to 10 (Fig. S9A) and the following parameters: iter.max = 10, nstart = 25. k = 4 was selected as the best compromise between parsimony (based on the diminishing decrease in the sum of squared errors) and interpretability (cluster visualization in Fig. 1F). Centroids reflecting the central selection coefficients of mutants from each cluster are visualized in Fig. S9B and provided in Table S5. To estimate relative enrichment of mutants from specific clusters at the sites of interest, we organized the data in separate contingency tables indicating whether a particular mutant was assigned or not to that site and its corresponding cluster. We calculated the log2 fold ratio of observed vs expected counts of mutants from each cluster at each site (Fig. S9C).

##### 8.3.2.- Random forest regressor

A random forest regressor was trained to identify the features that better explain variation in expression level dependent **Δ***s*. Explanatory variables included in the model were:

- Relative solvent accessibility (RSA): Solvent accessibility was calculated using DSSP version

2.2.1 (*54*) on the biological assembly of DfrB1 (PDB: 2RK1) (*7*). RSA was then obtained by dividing the solvent accessibility of each residue by the maximum solvent accessibility of that residue as described by Miller, et al. (*55*).

- Biophysical effects of mutations: FoldX version 5.0 (*56*, *57*) was used with the MutateX wrapper

(*58*) to simulate all possible mutations on the biological assembly of DfrB1 (PDB: 2RK1). Effects of mutations on subunit stability and binding affinity at the dimerization and tetramerization interfaces were measured.

- Differences in amino acid scores: 57 amino acid indexes were downloaded from the ProtScale database (*21*) on January 21st, 2019, as well as the index on propensity of each amino acid to participate in protein-protein interaction interfaces (*22*). For each mutation, we calculated the differences in index scores between the mutant residue and the WT residue.

All the explanatory variables were divided by their maximum values to scale them between -1 and 1. The random forest model was trained using the sklearn Python library (*59*, *60*) with 80 % of the data and tested with the remaining 20 %. We used the Python rfpimp library (https://github.com/parrt/random-forest-importances) to estimate relative importances of each variable by calculating the decrease in *R*^2^ of the predictions if the values of a particular feature are permuted (Fig. S16A). Due to the high degree of collinearity in our set of explanatory variables, retraining the random forest with different seeds can suggest a different set of top variables. We introduced a random variable *N(0, 1)* as an internal control in the training set to identify which variables have significant contributions to the model. We selected the variables that have a higher relative importance than the random variable and retrained the random forest to obtain the final model. Finally, we evaluated the relative importance of the variables in the final model by comparing the decrease in *R*^2^ when the final model is retrained without that variable (Fig. S16D).

#### 8.4.- Statistical analyses

##### 8.4.1.- ANOVAs

We used two different types of ANOVAs to characterize expression-dependent changes in fitness effects of coding mutations. First, we performed an ANOVA for each mutation considering all of its replicates at all the expression levels to identify mutants whose fitness effects were significantly affected by changes in gene expression. To correct for multiple hypothesis testing, we applied the Benjamini Hochberg correction with a false discovery rate of 0.05 using the FDRestimation R package (Table S4) (*61*). Then, we did an ANOVA on ranks with all replicates of all mutations at all the expression levels by applying the Aligned Rank Transform from the ARTool R package (*62*, *63*) to estimate the contribution of the interaction between expression level and coding mutants to fitness relative to their separate contributions (Table S6).

##### 8.4.2.- Generalized linear models (GLM)

We fitted GLMs to analyze the contributions of destabilizing effects of mutations to the measured selection coefficients (Table S8). This analysis was done separately for the data with TMP and without TMP using the glm function from the R base stats package (*53*). The formula for the GLMs used expression level, destabilizing effects (**ΔΔ***G* on subunit stability, **ΔΔ***G* on binding affinity at the dimerization interface, **ΔΔ***G* on binding affinity at the tetramerization interface), and terms for the interactions between expression level and each of the destabilizing effects.

## Supplementary Figures

**Fig. S1.**
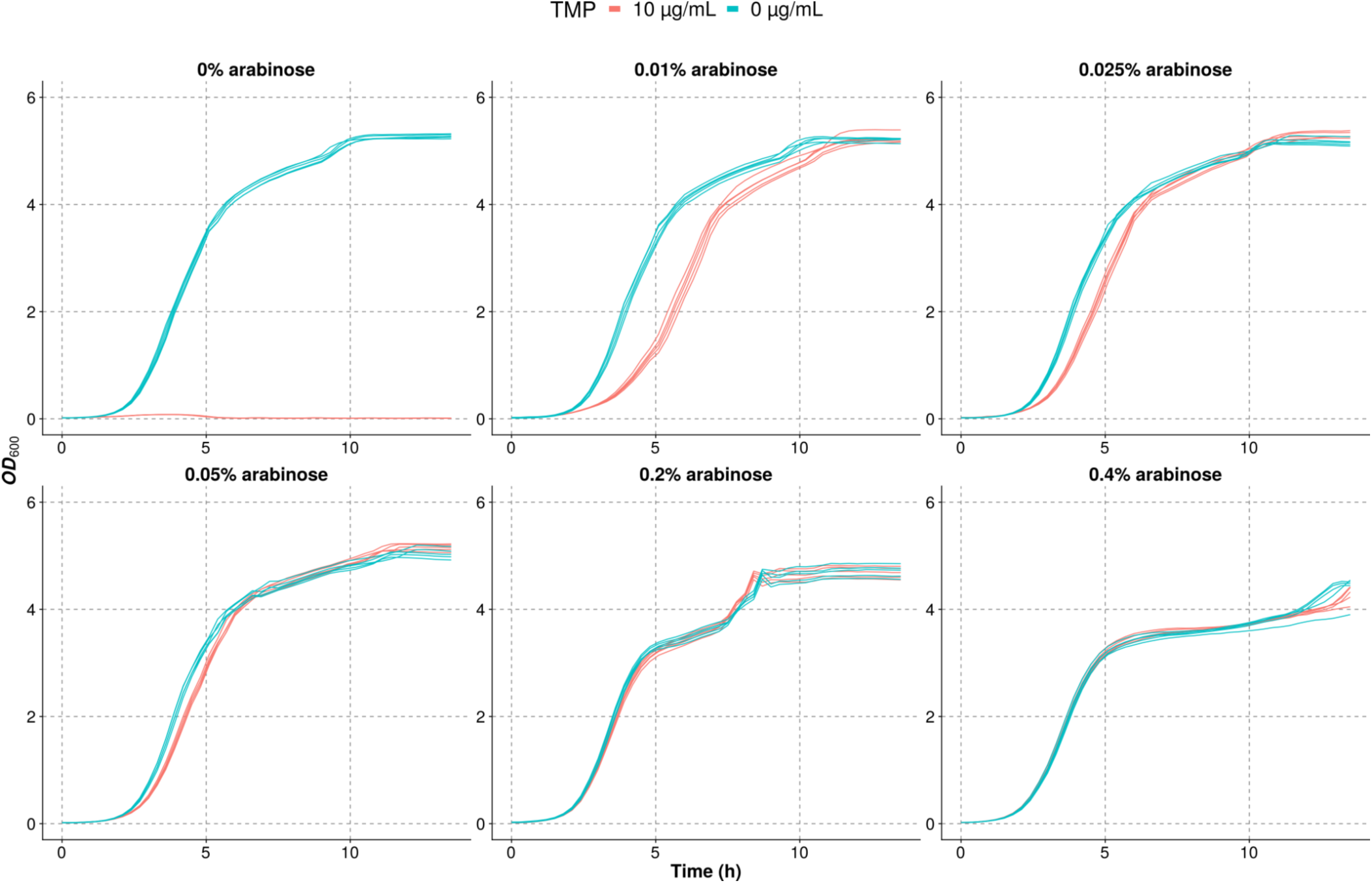
Dose-dependent growth recovery upon DfrB1 induction. Cells expressing WT DfrB1 were grown in LB medium with and without trimethoprim (TMP) and at different concentrations of arabinose indicated above each panel.

**Fig. S2.**
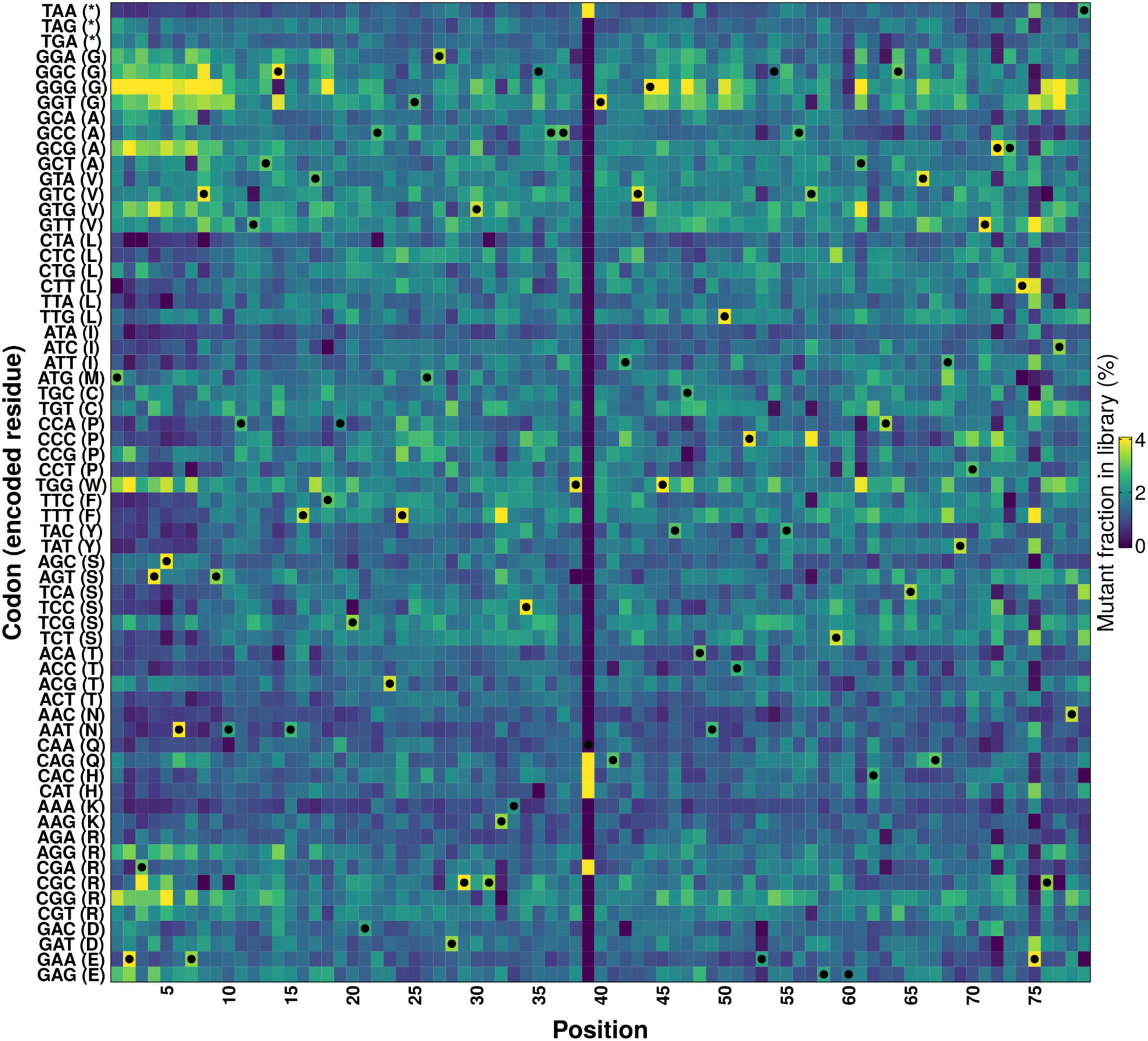
Quality control of the DfrB1 Deep Mutational Scanning (DMS) library. The libraries corresponding to each position were pooled and sequenced on a MiSeq PE300 (268 671 reads) to verify the coverage of the various codons across positions. A single position was not covered and was repeated and added to the final library. The color scale represents the percentage of reads that mapped to a mutant out of the library for that position. WT codons are labeled with dots.

**Fig. S3.**
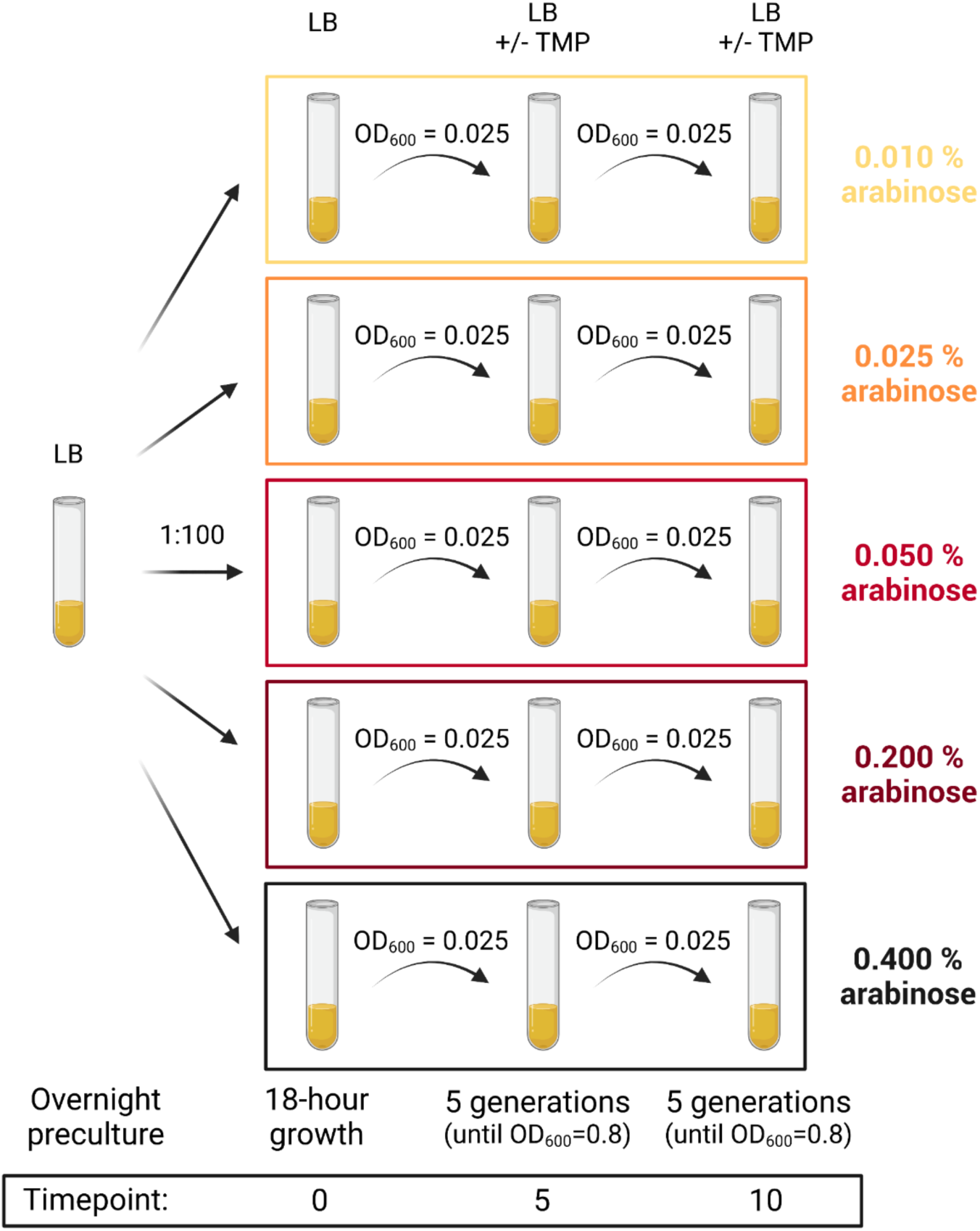
Experimental setup. Samples for the bulk competition experiments were taken from an overnight preculture of cells transformed with the DMS library. Cells were grown for five generations in LB medium with different concentrations of arabinose and with or without trimethoprim (TMP) until they reached an optical density of 0.8. They were then diluted again and grown for five more generations. DNA was extracted and sequenced at t = 0 and t = 10 generations. This experiment was repeated twice (see Table S1 for details).

**Fig. S4.**
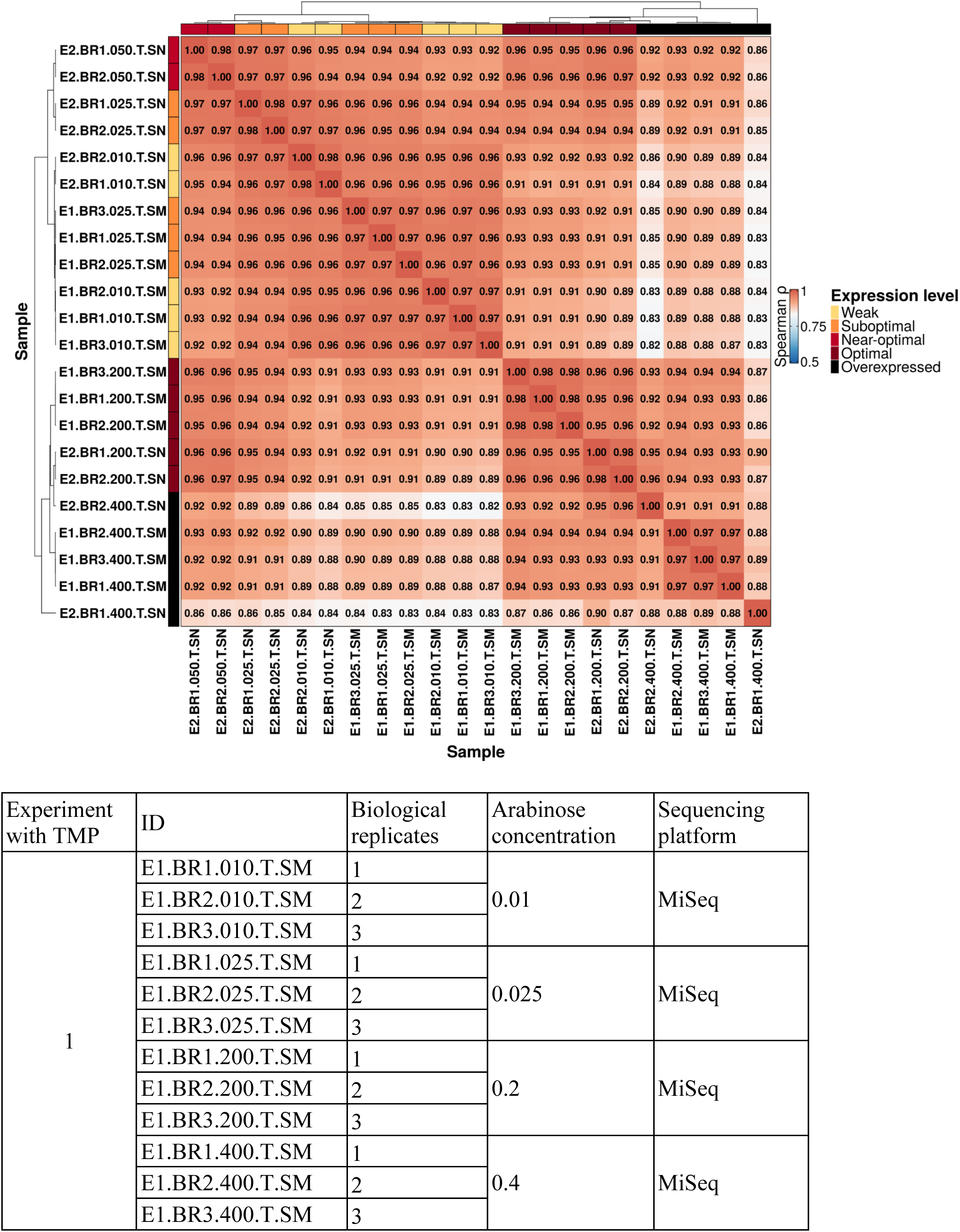
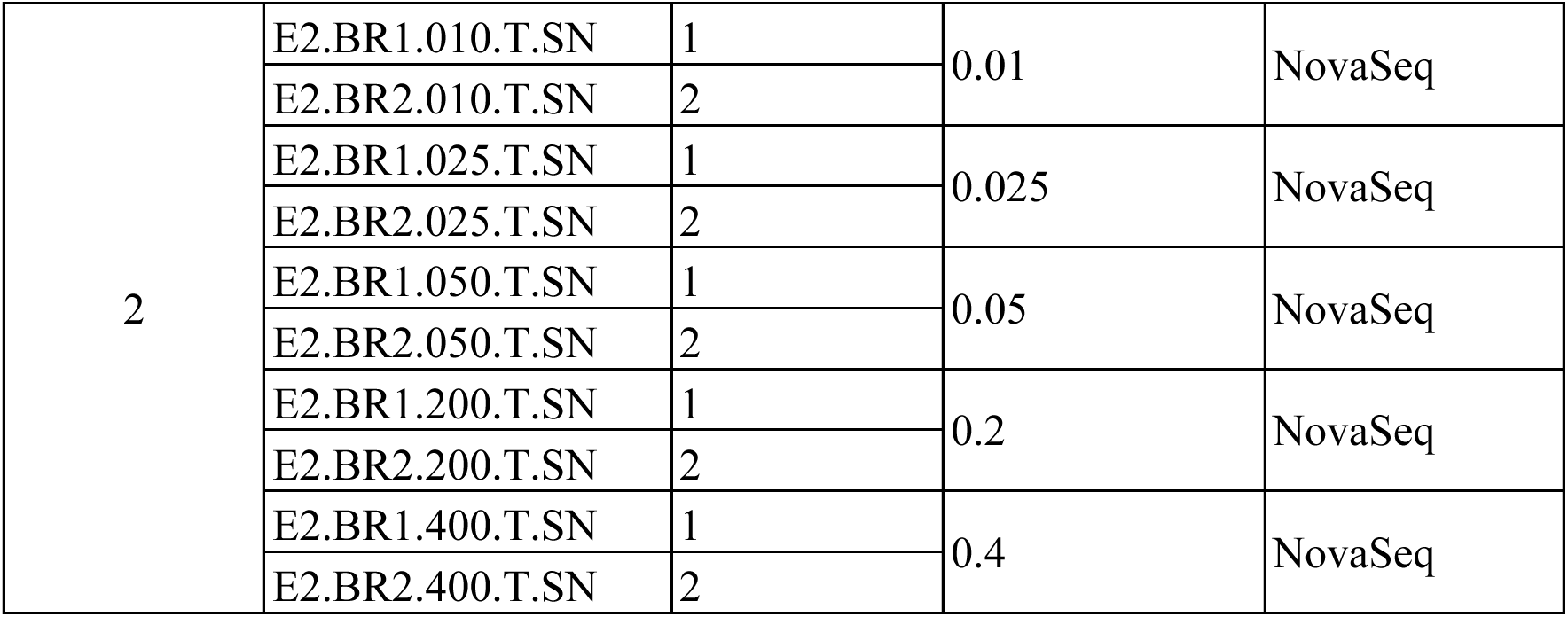
Replicates of the bulk competition experiment with selection for DfrB1 activity (with TMP) correlate strongly. Spearman correlation between selection coefficients was estimated for different replicates of the experiment with selection for DfrB1 activity. Samples were named according to the table (bottom) extracted from Table S1.

**Fig. S5.**
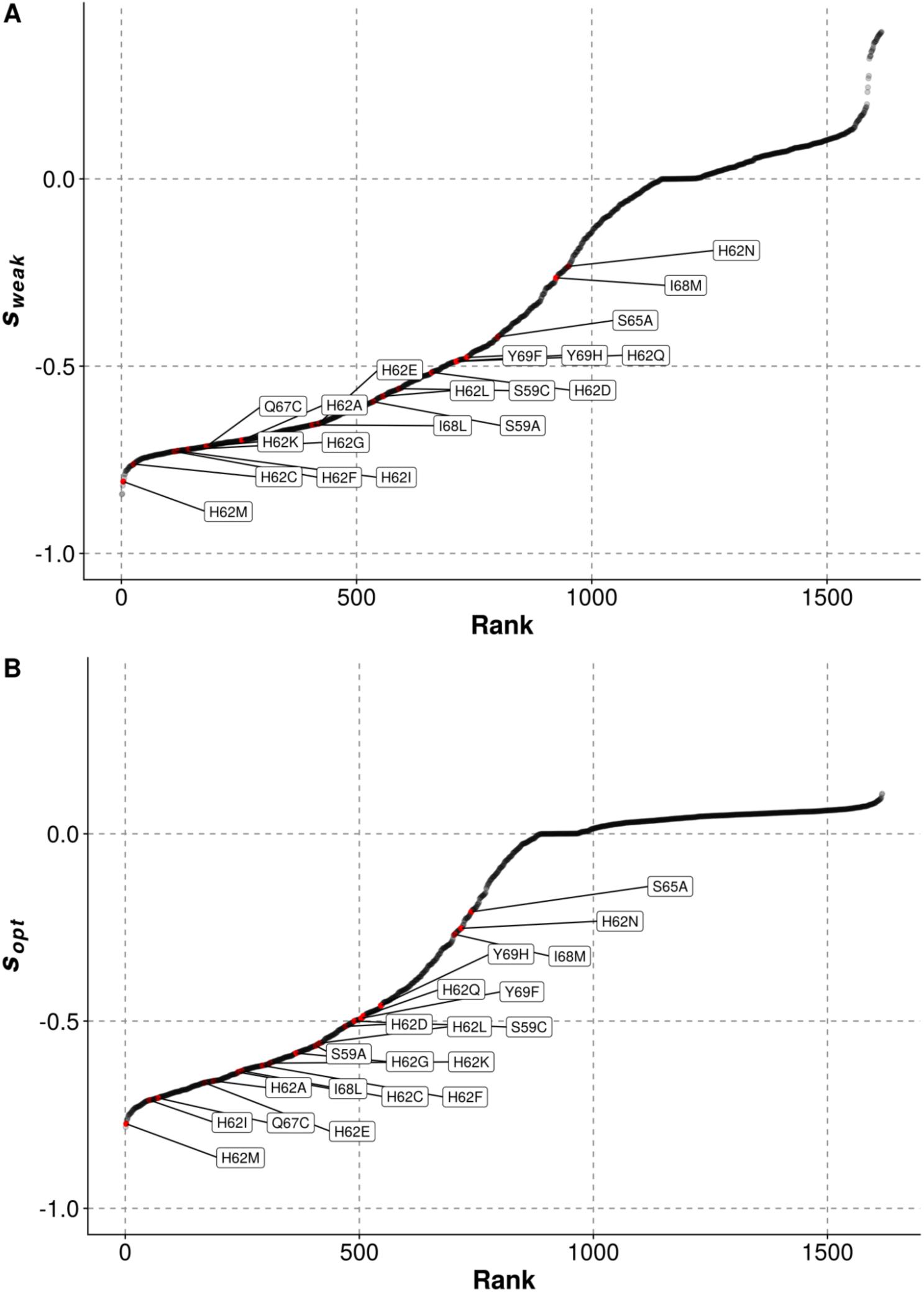
Selection coefficients recapitulate previously characterized poorly active mutants. Ranks and selection coefficients in our bulk competition assay for deleterious mutants constructed by Dam et al. (*64*) and Strader et al. (*16*) at a weak expression level for DfrB1 **(A)** and at the optimal expression level (B). All mutants have negative *s*, as expected.

**Fig. S6.**
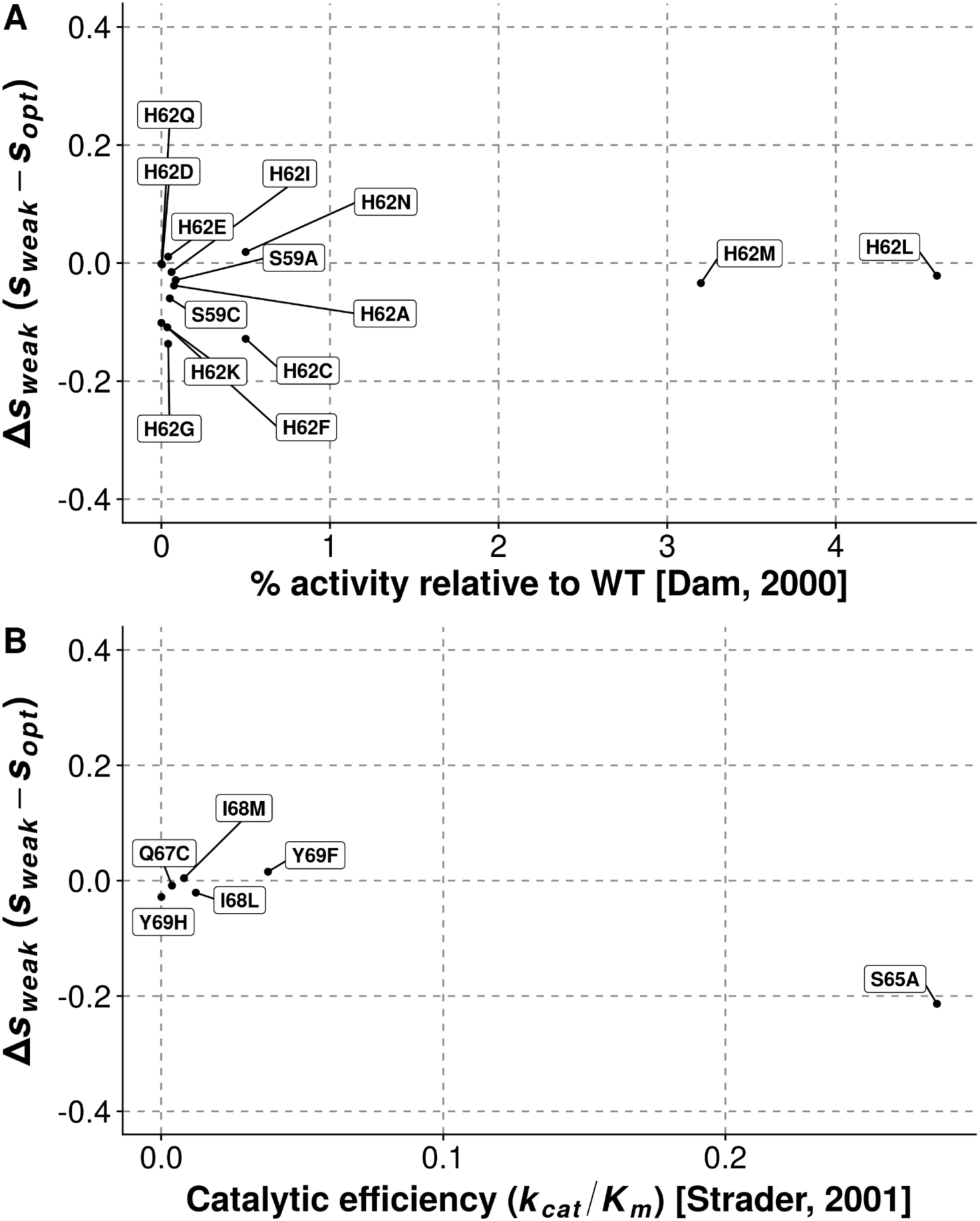
The fitness effects of only some of the low activity mutants are improved at optimal expression. (**A**) Differences in selection coefficients observed at weak expression and optimal expression on mutants at the tetramerization interface from Dam et al. (*64*) with respect to their measured activity relative to WT. (B) Differences in selection coefficients observed at weak expression and optimal expression on active site mutants from Strader et al. (*16*) with respect to their measured catalytic efficiency.

**Fig. S7.**
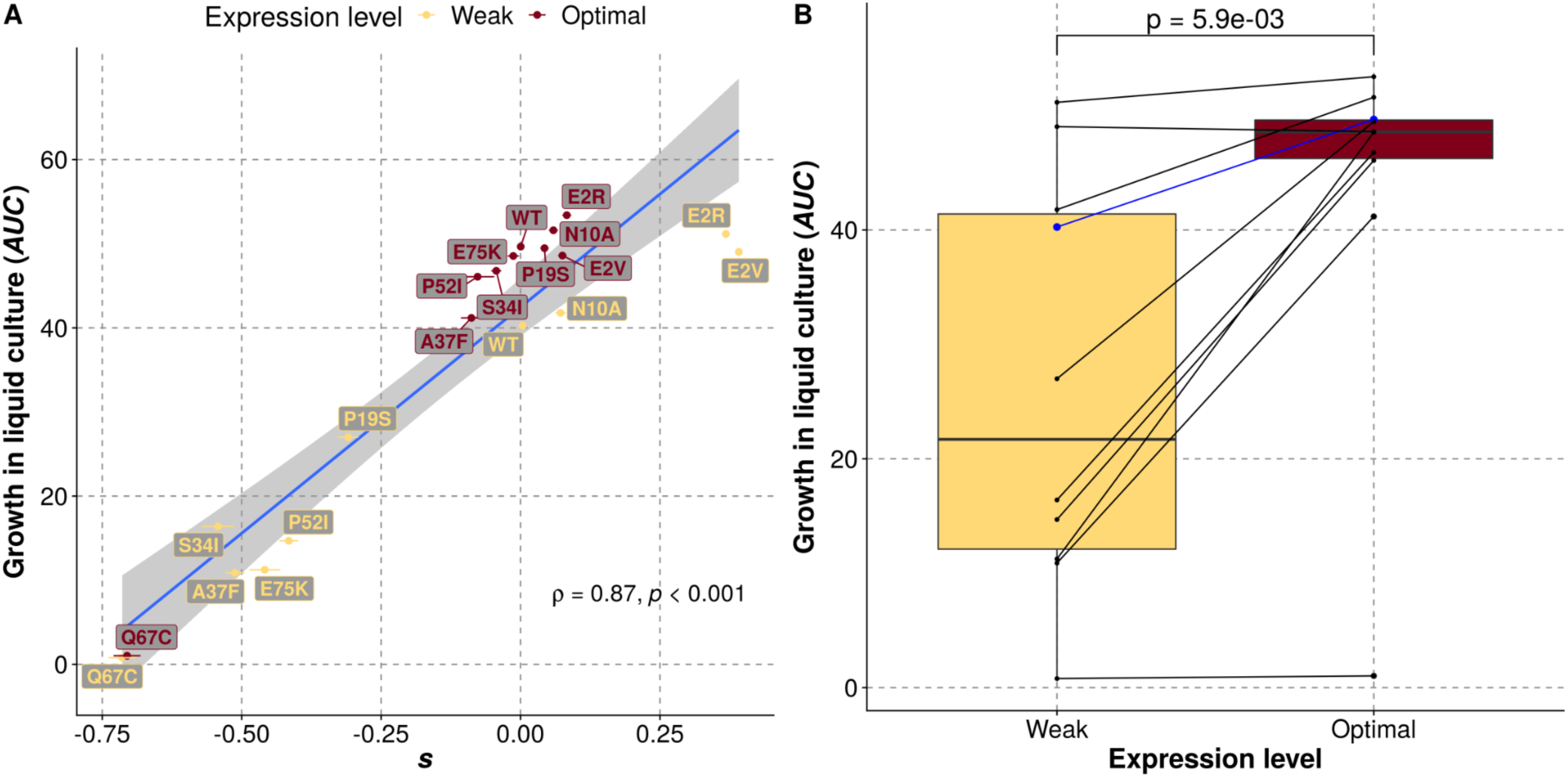
The effects of mutations measured by bulk competition are recapitulated with individual growth measurements. Mutants with different magnitudes of changes in fitness effects across concentrations of arabinose were selected for validation in individual growth experiments. Growth was measured as the average of the area under the curve for two replicates per mutant at each of two expression levels. (**A**) Growth of individual mutants correlates strongly with DMS normalized fitness scores. The corresponding mutants and the WT are indicated next to each data point. (B) Expression-level dependent differences in growth rate for the validated mutants. WT is indicated in blue. P-values were calculated using Wilcoxon’s test for differences in means of paired samples.

**Fig. S8.**
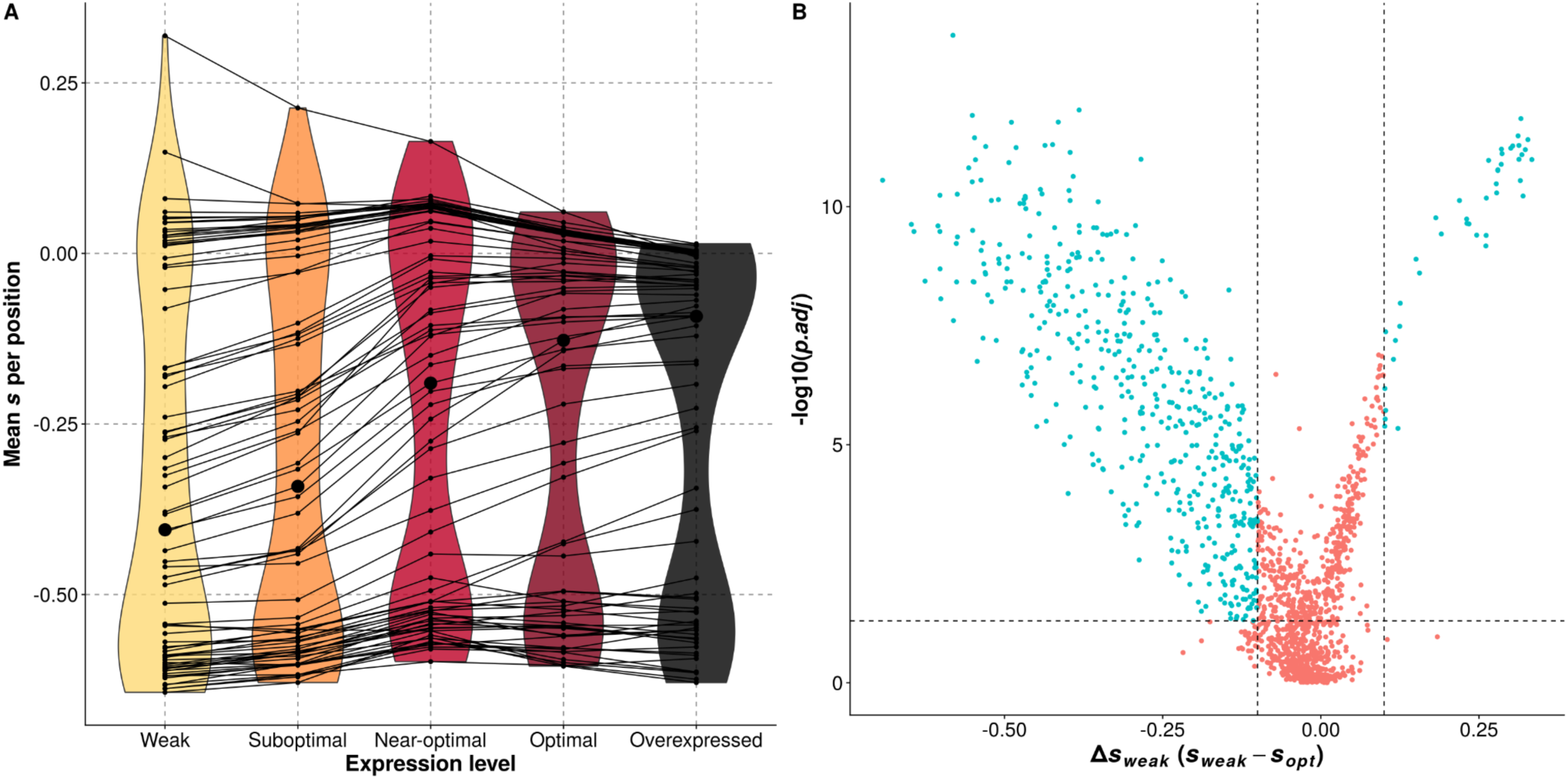
Significance and magnitude of effects of expression level on the fitness of mutants. (**A**) Averages of selection coefficients observed at each position of the DfrB1 sequence. The larger dot indicates the position with the median effect for each expression level. (B) For each mutant, we performed an ANOVA and corrected for multiple hypothesis testing by applying the Benjamini-Hochberg correction with a false discovery rate of 0.05 using the FDR estimation package (*61*). The horizontal dashed line indicates the cutoff for significance of adjusted p-values (p < 0.05) and the vertical lines indicate an arbitrary threshold for magnitude of changes in selection coefficients (greater than 0.1) due to expression.

**Fig. S9.**
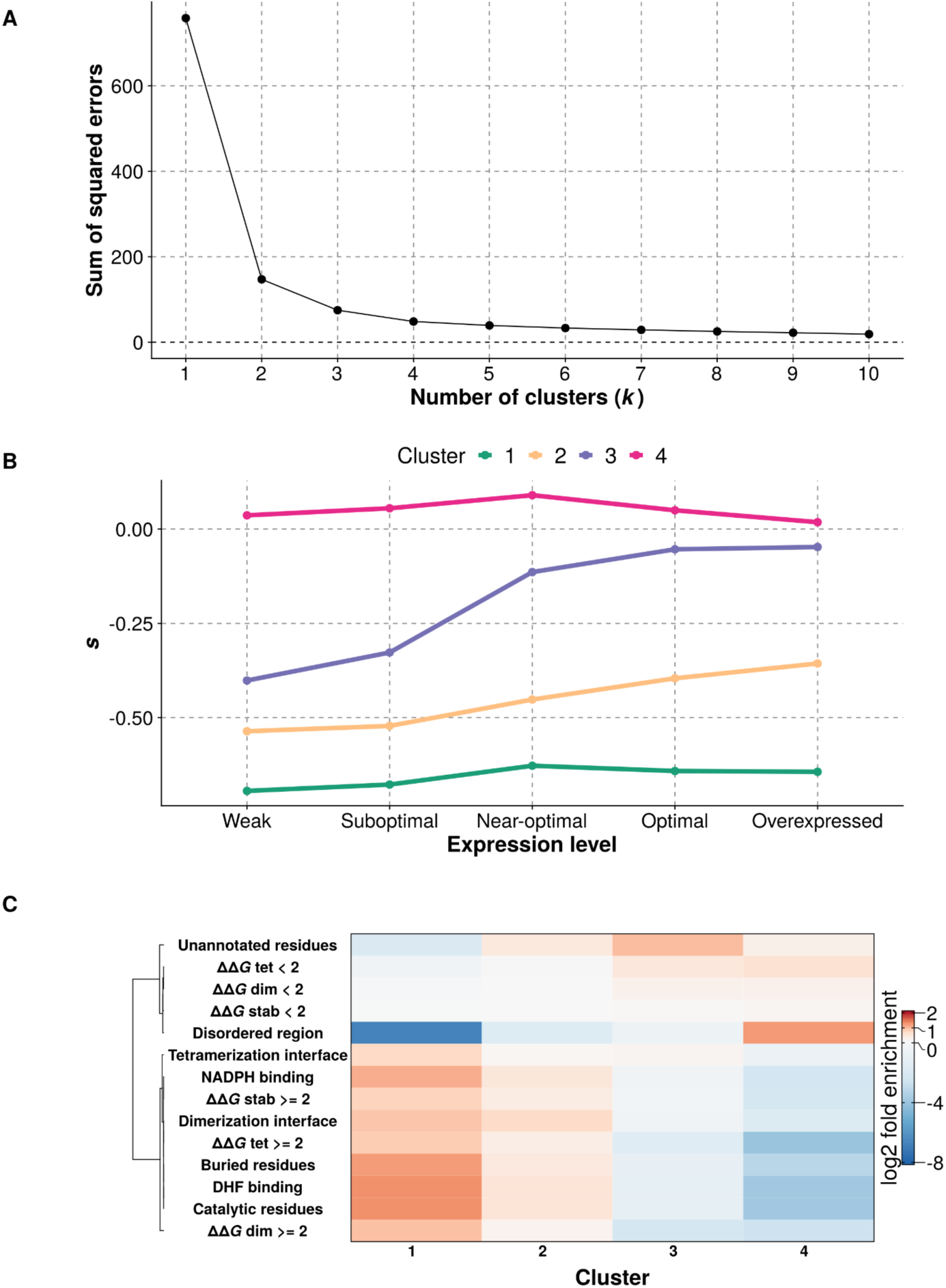
K-means clusters capture the main patterns of expression-dependent changes in fitness effects. (**A**) Sum of squared errors as a function of the numbers used in k-means clustering. (B) Selection coefficients at each expression level for the centroids of the four clusters. (C) Relative enrichment of each of the four clusters with mutants at particular protein sites and with specific destabilizing effects.

**Fig. S10.**
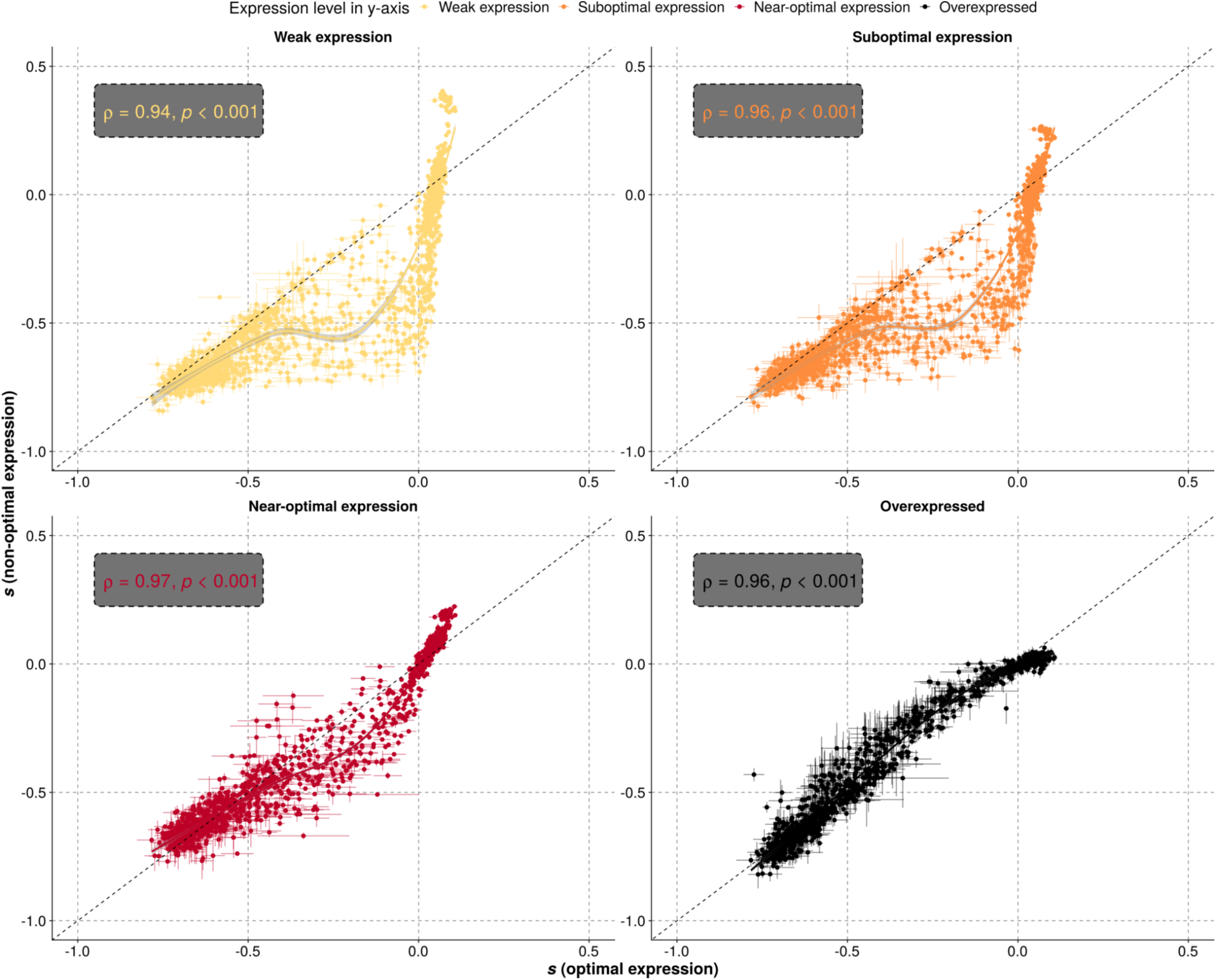
Fitness effects of mutations are dependent on expression level. The fitness effects of mutants at optimal expression are shown on the x-axis and compared to the fitness effects of mutants at lower or higher expression level. Same data as in Fig 1E and Fig. 2. The solid line indicates a LOESS regression. Horizontal error bars indicate the standard error of the mean observed across biological replicates at optimal expression level. Vertical error bars indicate the standard error of the mean observed across biological replicates at non-optimal expression levels.

**Fig. S11.**
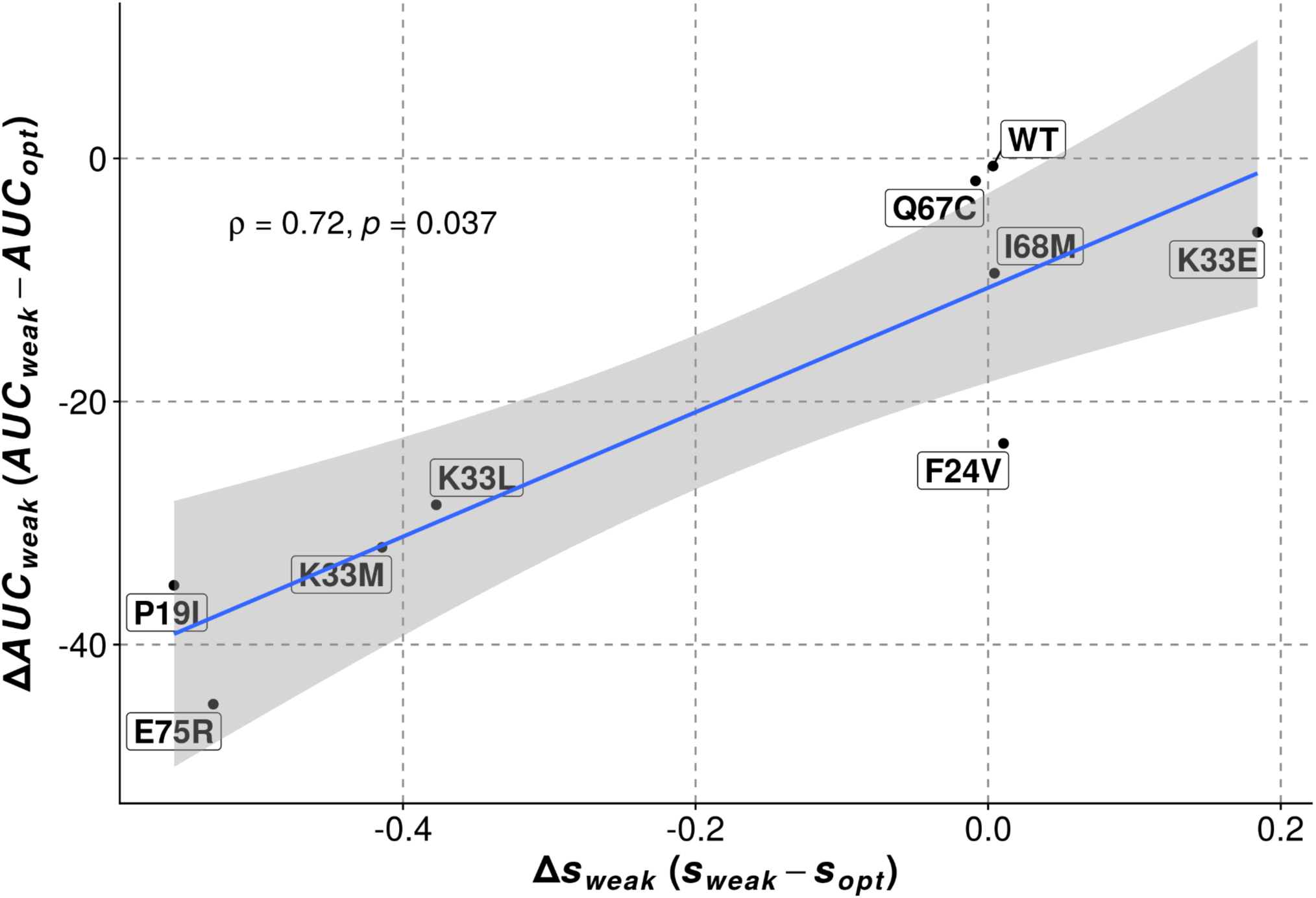
Liquid cultures recapitulate well the expression level dependent differences in fitness effects in the bulk competition experiment. **Δ***AUC_weak_* was calculated as the differences in the area under the curve observed for growth in liquid cultures at weak and optimal expression levels (*AUC_weak_* - *AUC_opt_*). **Δ***s_weak_* was calculated as the differences in selection coefficients observed for each mutant in the bulk competition experiments at weak and optimal expression levels (*s_weak_* - *s_opt_*).

**Fig. S12.**
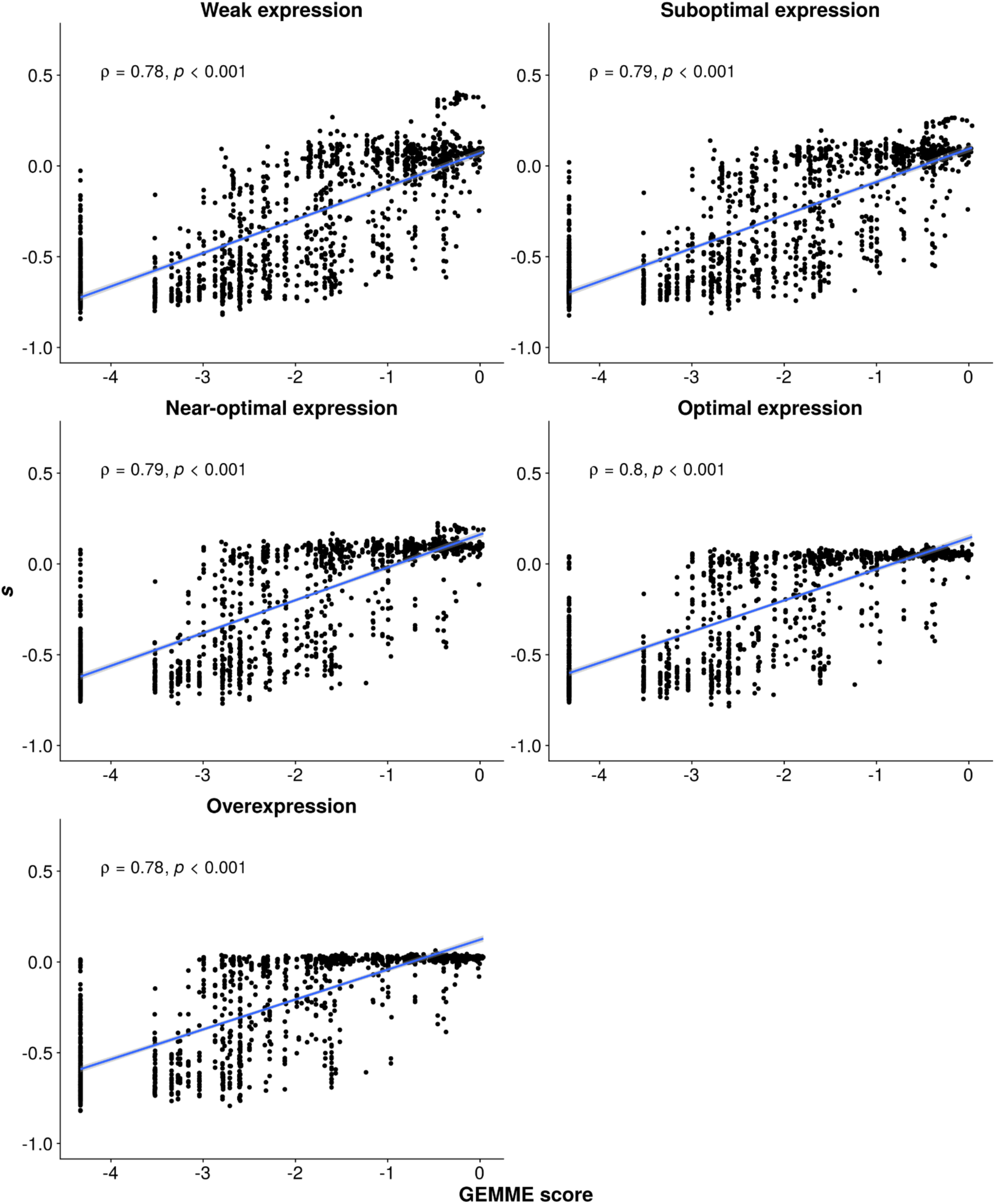
Selection coefficients measured in the DMS bulk competition experiment with TMP correlate well with fitness effects deduced variation observed in natural sequences. We used GEMME (*11*) to predict mutational effects based on patterns of conservation and substitution. GEMME scores (0 for residues observed as often as the WT DfrB1 residue at that position and negative for residues observed less often) correlate well with the effects measured in the bulk competition assay with the mutant libraries. These analyses show that the mutational effects measured experimentally reflect the selective pressures these homologous sequences experience in nature.

**Fig. S13.**
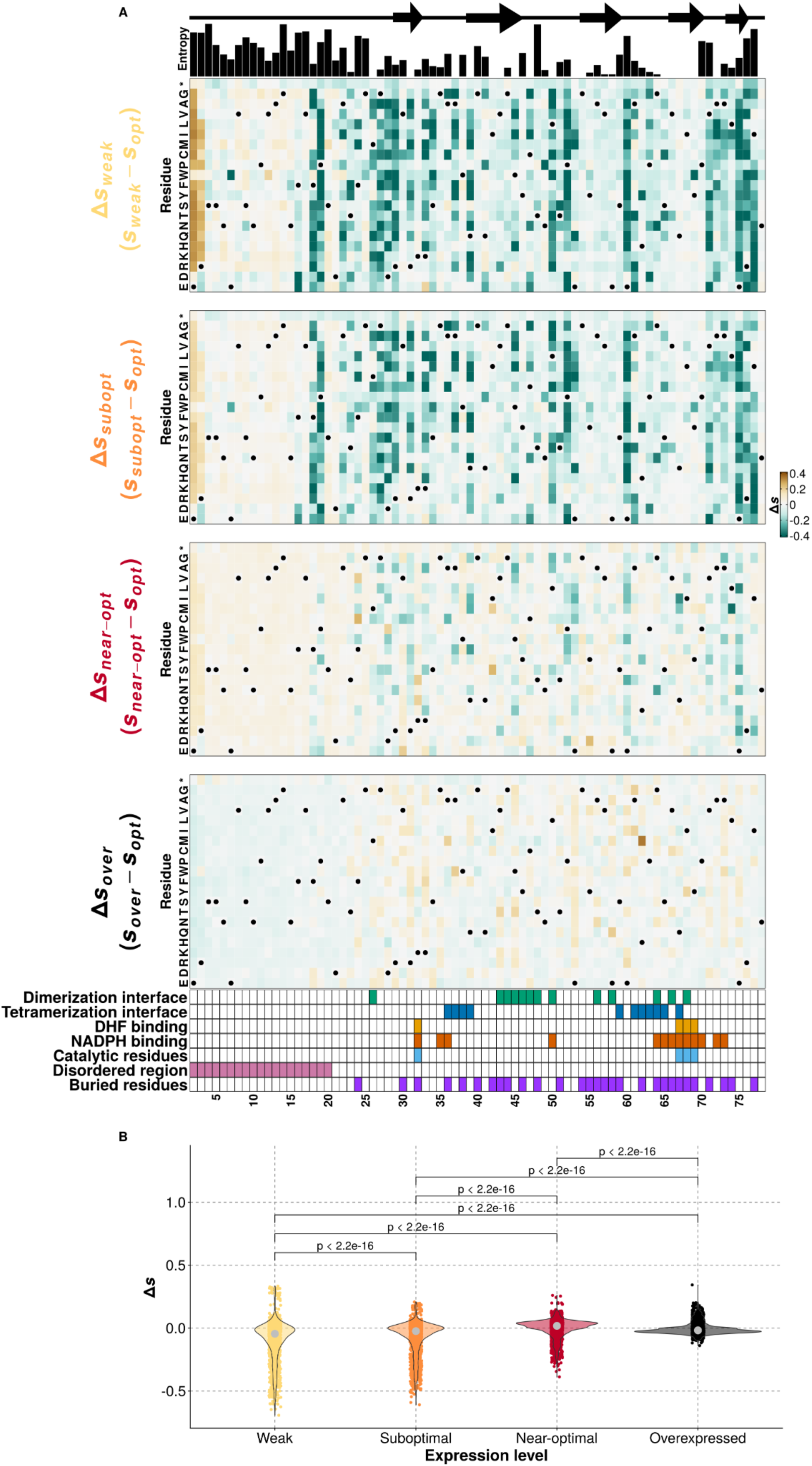
Expression-dependent differences in fitness effects become weaker as expression level approaches the optimum. (**A-B**) Difference between the selection coefficients of an amino acid substitution at one of the non-optimal expression levels and at the optimal expression for the WT (**Δ***s_non-opt_* = *s_non-opt_* - *s_opt_*). WT residues in panel A are labeled with dots. P-values in panel B were calculated using Wilcoxon’s test for differences in means of paired samples.

**Fig. S14.**
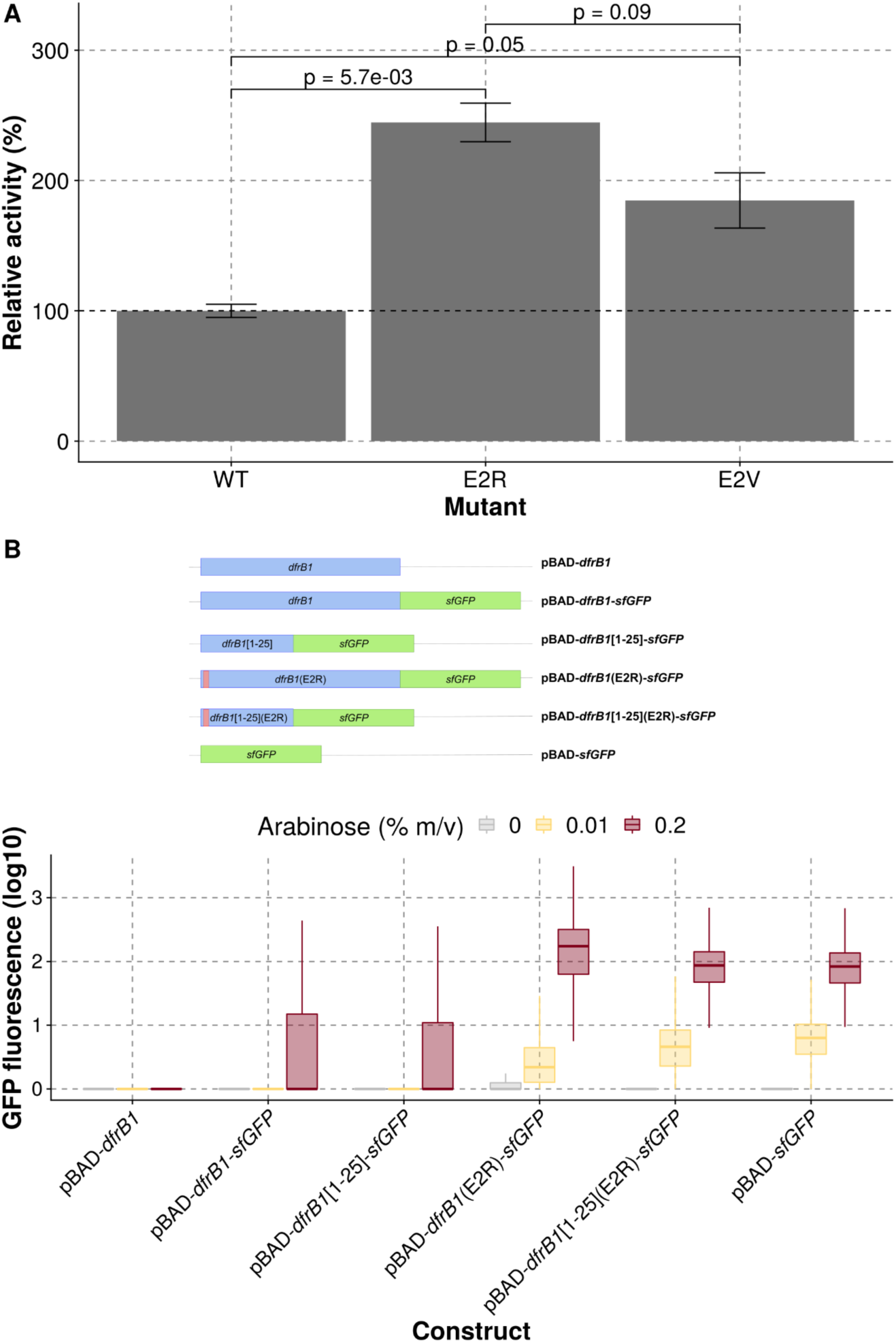
Mutants E2R and E2V have increased activity in cell lysates and expression level. (**A**) Comparison of relative activities observed for WT DfrB1 and mutants E2R and E2V. Bars indicate the means and error bars indicate the standard error of the mean of three replicates for each mutant. P-values were calculated with a t-test. (B) The fluorescence of GFP fused to several constructs of DfrB1 (top) was measured by flow cytometry (bottom). Constructs used include: 1) DfrB1 alone, 2) DfrB1 fused to GFP, 3) truncated DfrB1 (positions 1-25) fused to DfrB1, 4) E2R DfrB1 mutant fused to GFP, 5) E2R truncated DfrB1 mutant (positions 1-25) fused to GFP, 6) GFP alone.

**Fig. S15.**
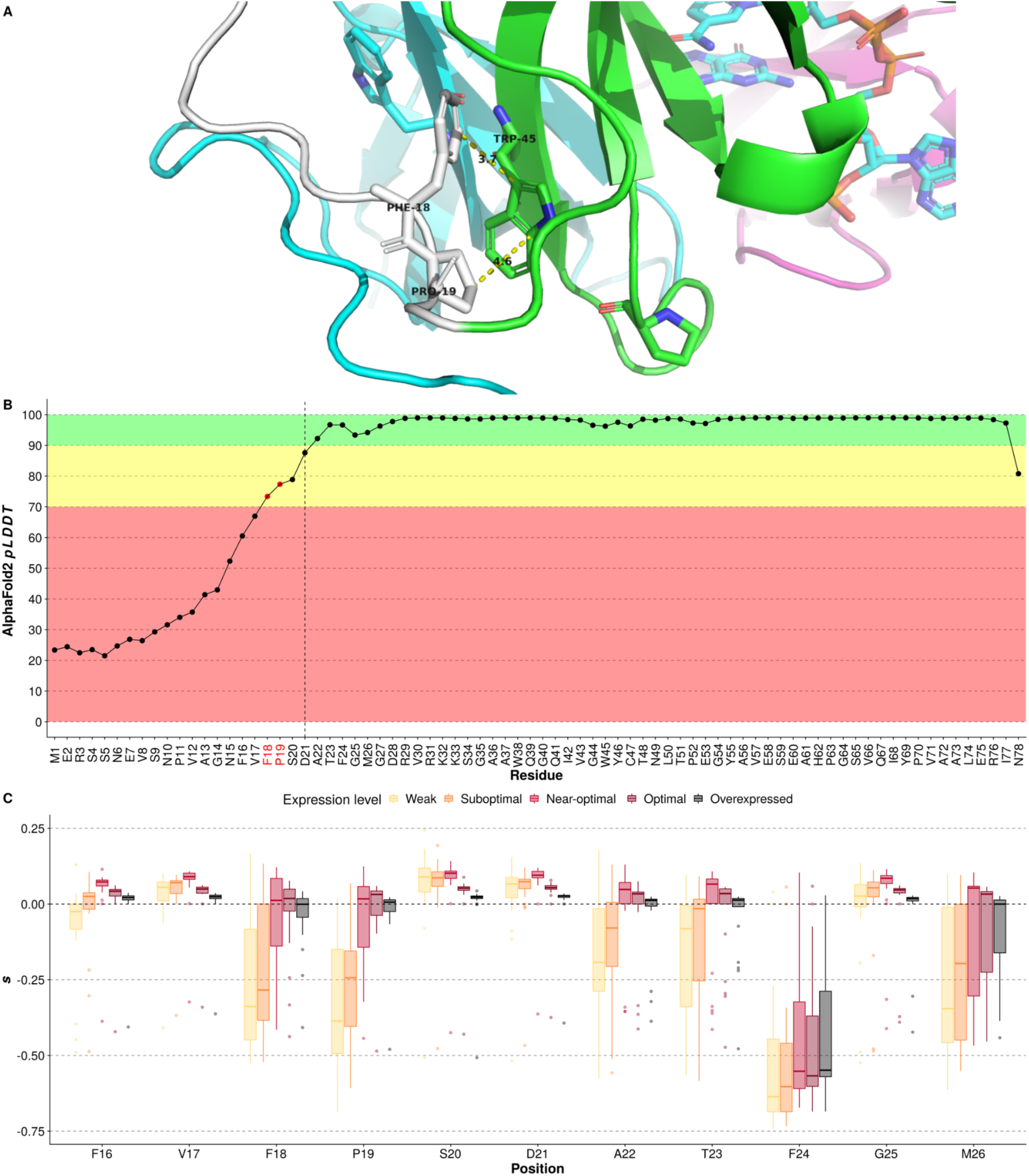
Predicted interactions of W45 with F18 and P19. (**A**) Residues F18 and P19 are predicted to interact with W45 in an AlphaFold2 predicted model. Residues F18, P19, and W45, as well as measurements are labeled. The disordered region of subunit A is shown in white, the rest of subunit A is shown in green, subunit B is shown in magenta, subunit C is shown in cyan. The structure was visualized with PyMOL (*65*). (B) *pLDDT* values assigned by AlphaFold2 to each of the predicted residues. Background colors indicate confidence levels: high confidence (*pLDDT* > 90, green), overall good backbone prediction (70 > *pLDDT* > 90, yellow), and low confidence (*pLDDT* < 70, red). Positions to the left of the dashed vertical line are not present in the crystal structure for DfrB1 (PDB: 2RK1). (C) Distributions of mutational effects for residues between positions 16 - 26 show that mutations at positions 18 and 19 are deleterious at low expression levels but can be masked at the optimal expression for the WT.

**Fig. S16.**
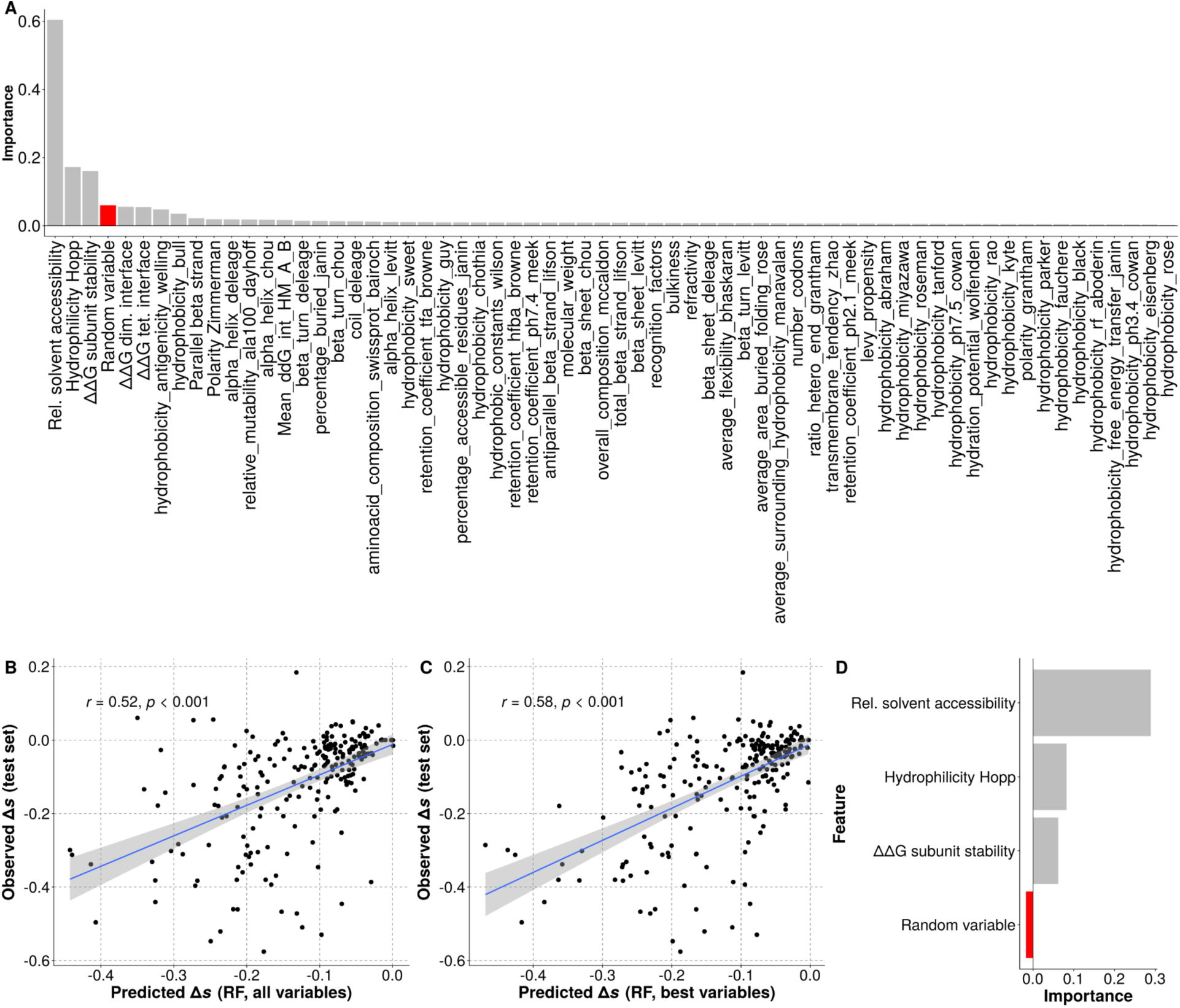
Identifying the most relevant features used to predict expression level dependent differences in selection coefficients. (**A**) Relative importance of all the features considered in the random forest model as calculated by the decrease in out-of-bag *R*^2^ when permuting the values of that feature. A random variable was included to identify the features that contribute the most to the model. Table S7 contains a list of all variables used in this analysis. (B) Comparison of observed **Δ***s_weak_* (*s_weak_* - *s_opt_*) versus predicted **Δ***s_weak_* using a random forest (RF) regressor with all the features for the test set (20 % of the total dataset). (C) Comparison of observed **Δ***s_weak_* (*s_weak_* - *s_opt_*) versus predicted **Δ***s_weak_* using an RF regressor with all the top three features for the test set (20 % of the total dataset). (D) Relative importance of the top features in the final model as calculated by the decrease in out-of-bag *R*^2^ when dropping one variable at a time and retraining the model.

**Fig. S17.**
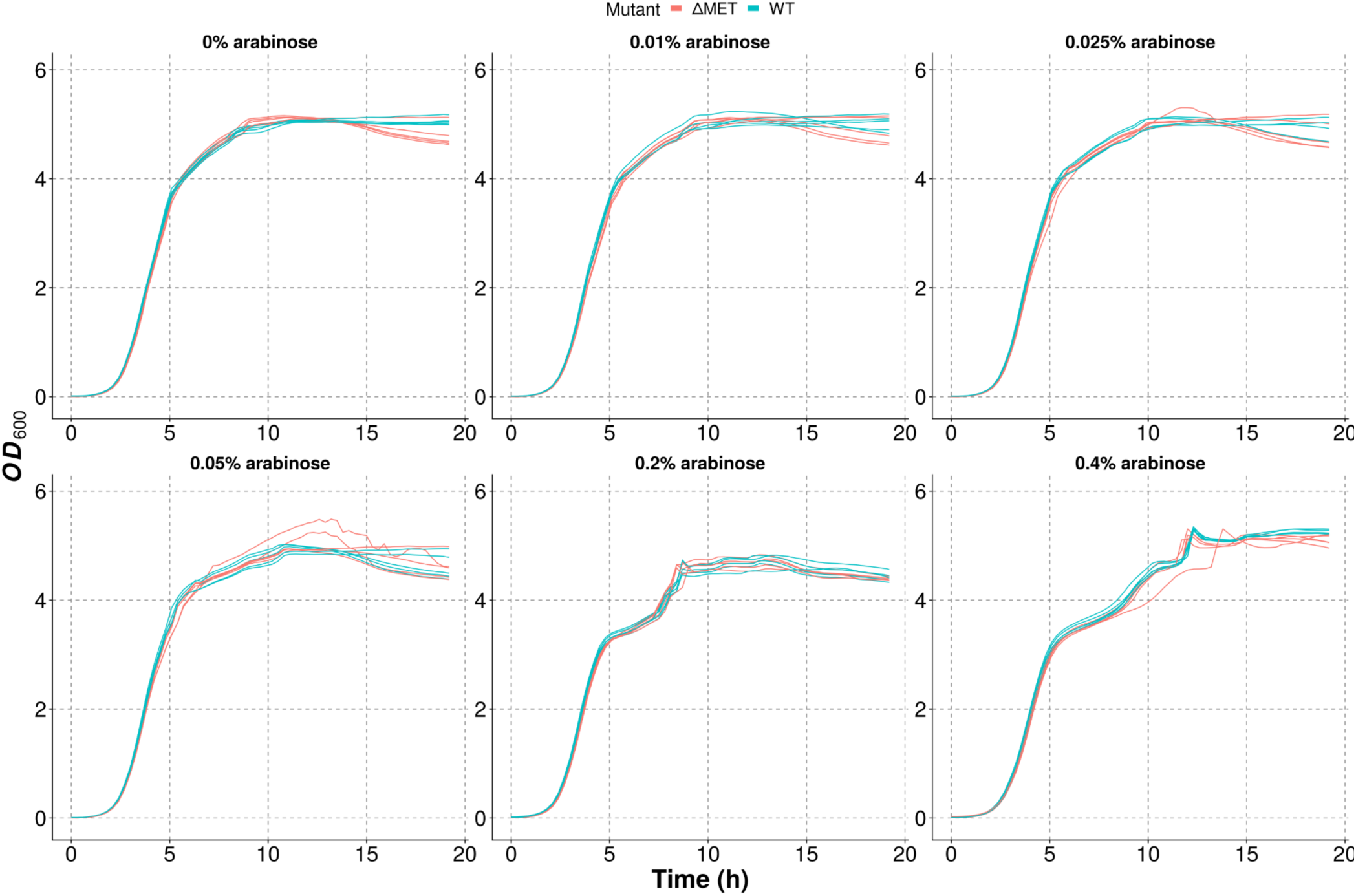
No clear toxic effects of expression of DfrB1. Growth curves for cells transformed with WT DfrB1 or with a DfrB1 mutant with a stop codon at position 1 (**Δ**MET) and grown in medium without TMP.

**Fig. S18.**
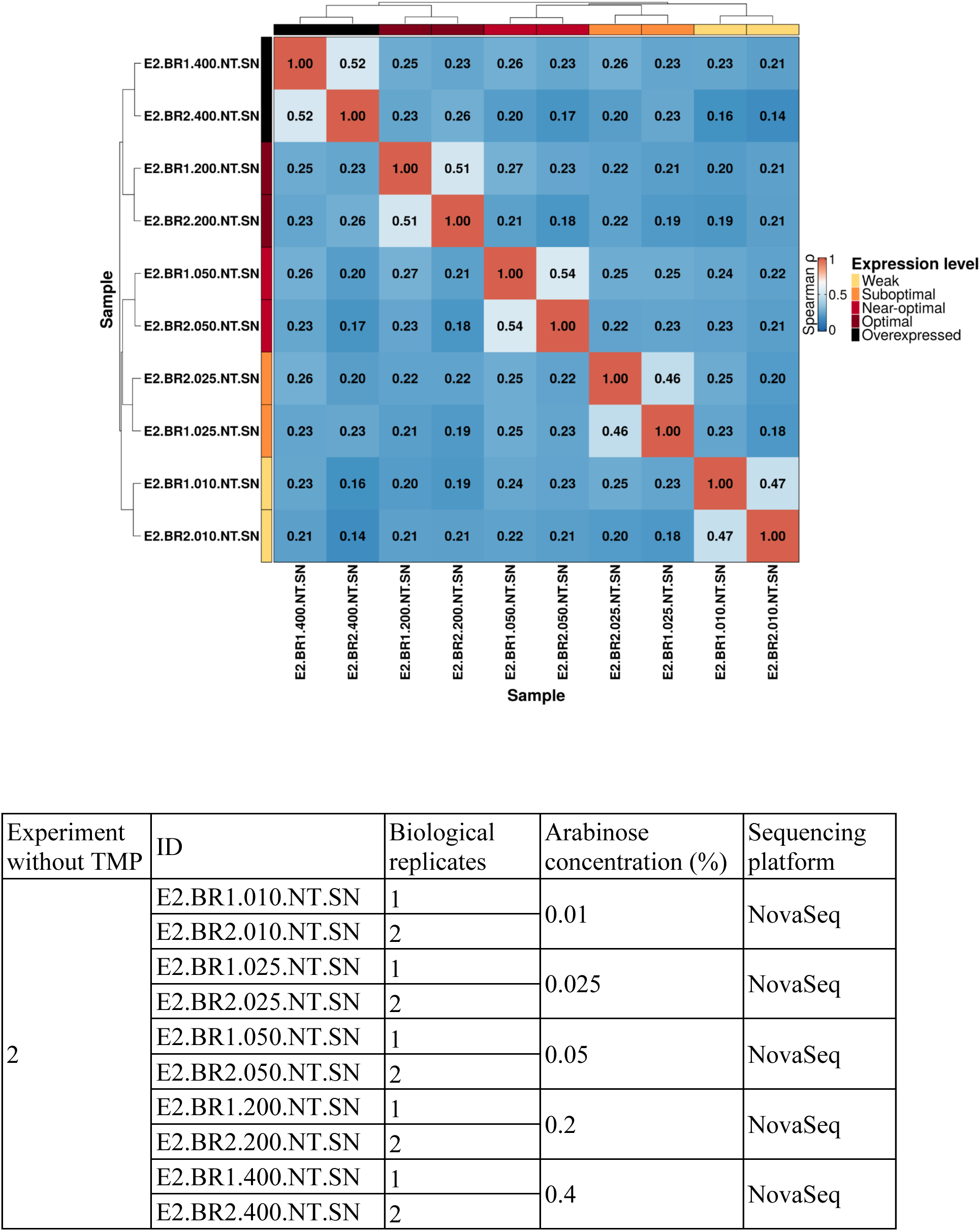
Low signal-to-noise ratio in selection coefficients in the absence of selection for DfrB1 activity (no TMP). Spearman correlation between selection coefficients was estimated for different replicates of the experiment without TMP. Samples were named according to the table (bottom) extracted from Table S1.

**Fig. S19.**
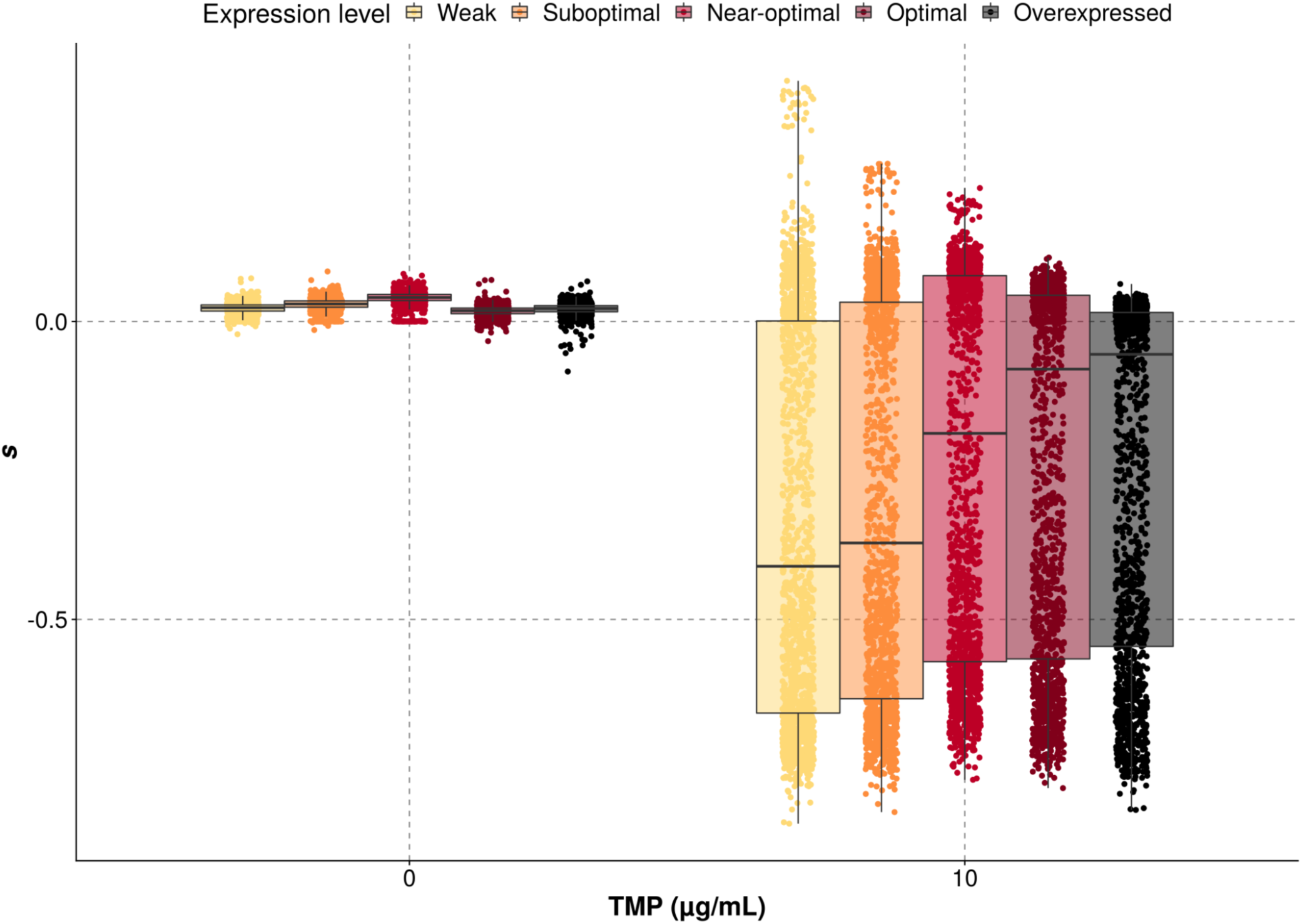
Overall distributions of selection coefficients in the experiments with and without selection for DfrB1 activity. The range of selection coefficients observed in the experiment with selection for DfrB1 activity was much broader than in the experiment without selection.

**Fig. S20.**
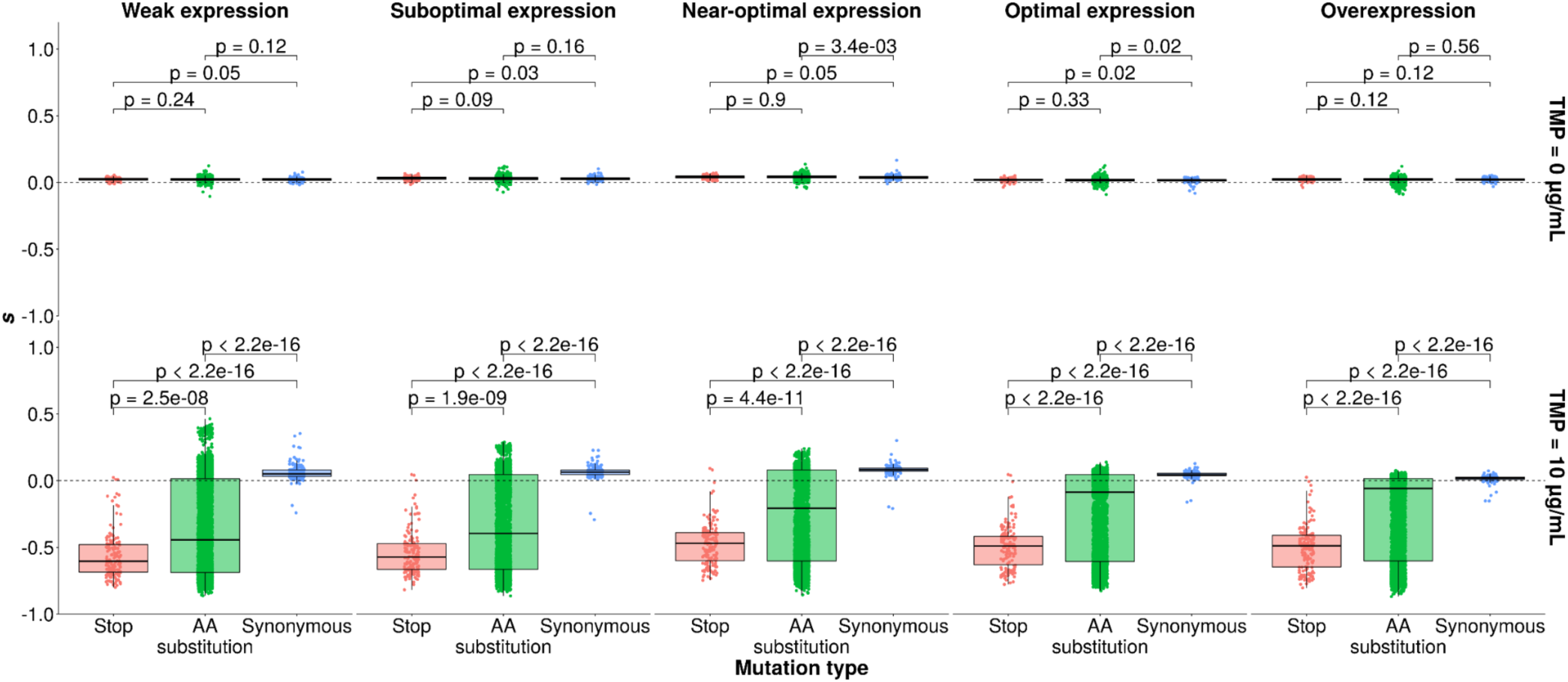
Distributions of selection coefficients for stop codons, amino acid substitutions, and synonymous mutants. Mutations introducing amino acid substitutions or stop codons are deleterious in the experiment with selection for DfrB1 activity (with TMP), but not in the experiment without selection for DfrB1 activity (without TMP). P-values were calculated using Wilcoxon tests.

**Fig. S21.**
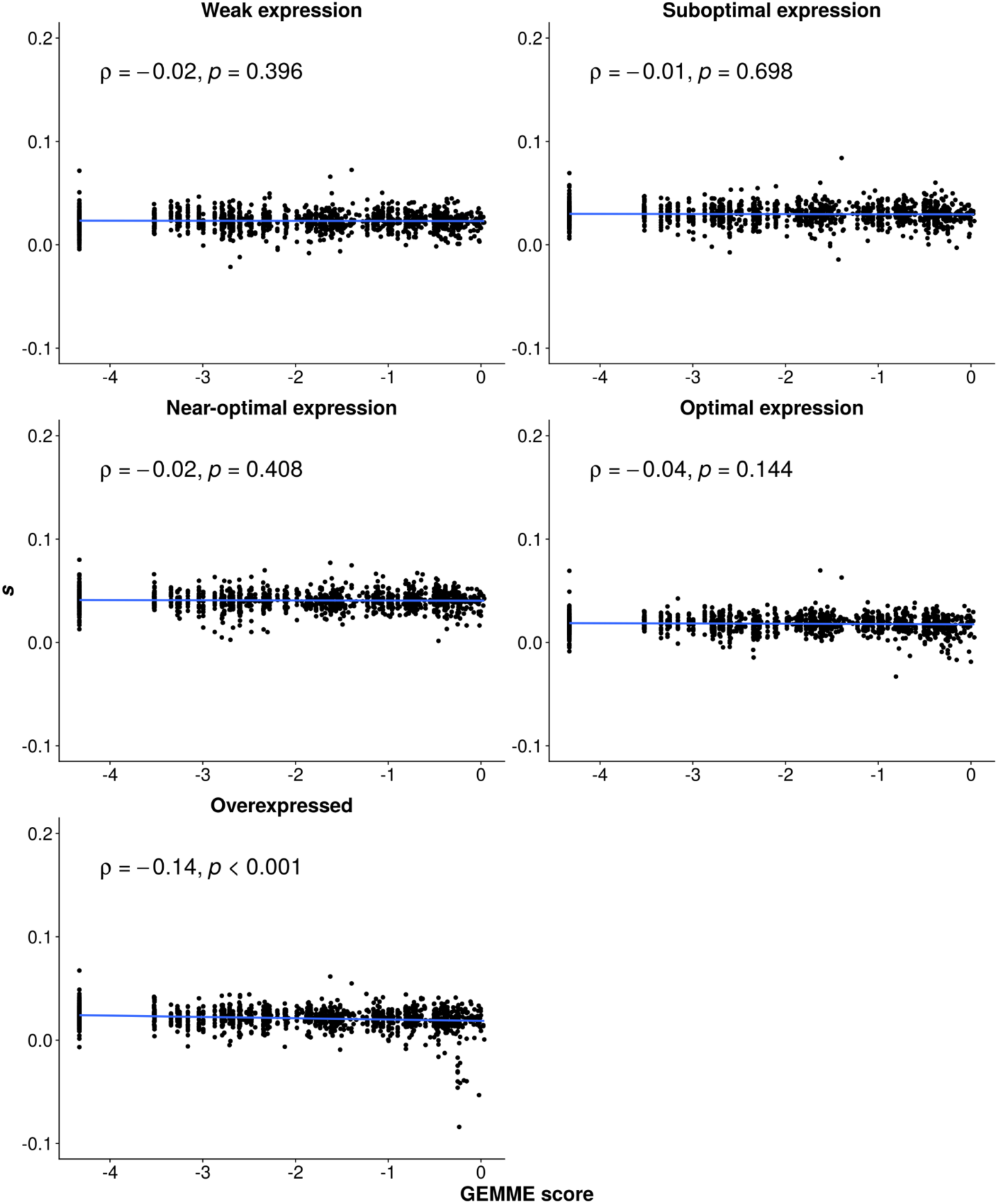
Fitness effects measured in the DMS bulk competition experiment without TMP do not correlate well with fitness effects deduced variation observed in natural sequences. The GEMME scores from Fig. S12 were used.

**Fig. S22.**
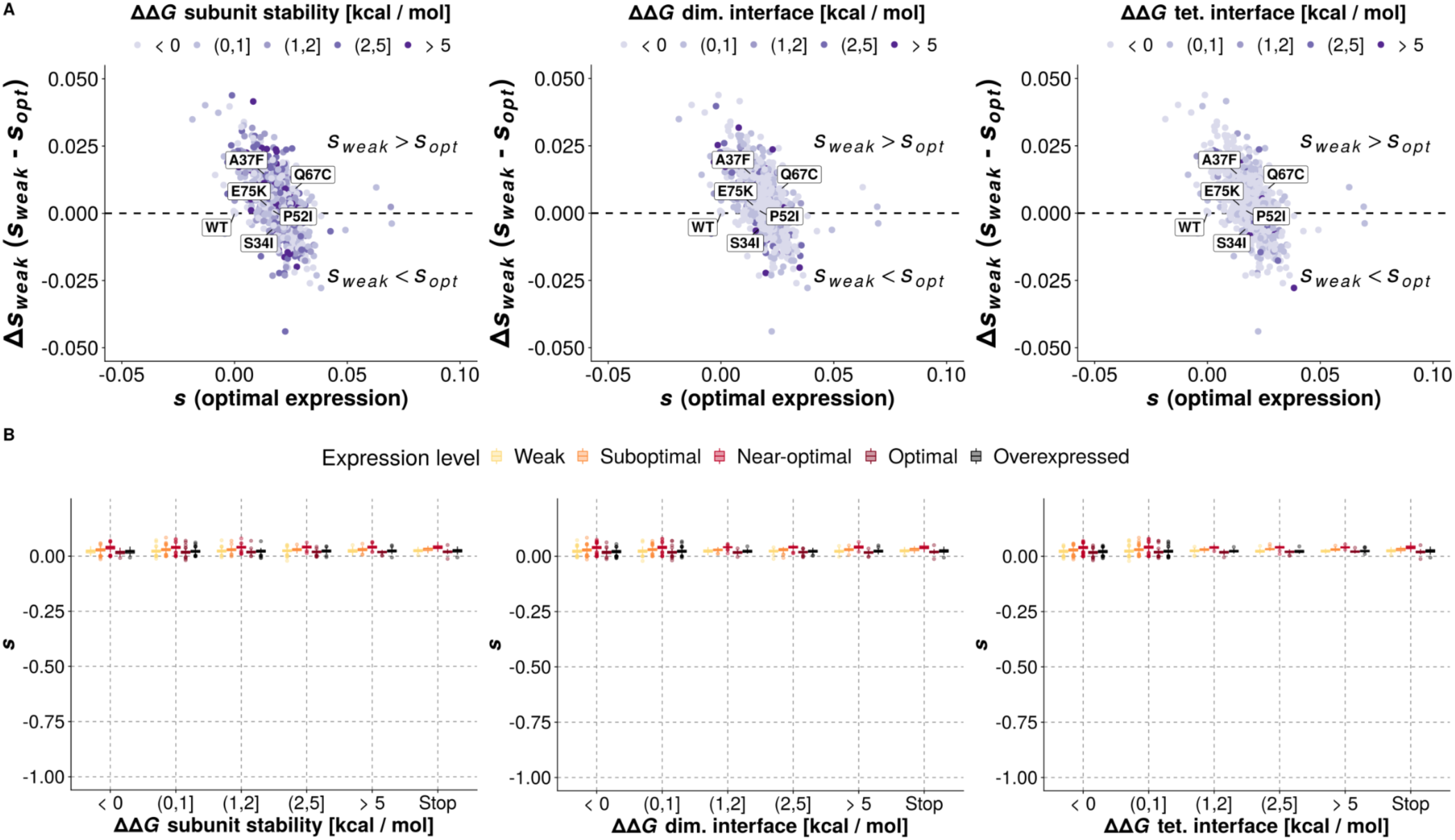
Protein destabilization by itself does not result in deleterious effects in the absence of selection for DfrB1 activity. (**A**) Landscape of selection coefficients measured at optimal expression (x-axis) and the expression-dependent change in fitness effects (y-axis) labeled with respect to mutational effects on subunit stability and binding affinity. (B) Distributions of measured selection coefficients for mutants with different effects on protein stability or binding affinity at the interfaces. Stop codons from the functional protein core (residues 30 - 70) are shown for reference. The scale in panel B is the same as in Fig. 4B for comparison with the experiment with selection for DfrB1 activity.

**Fig. S23.**
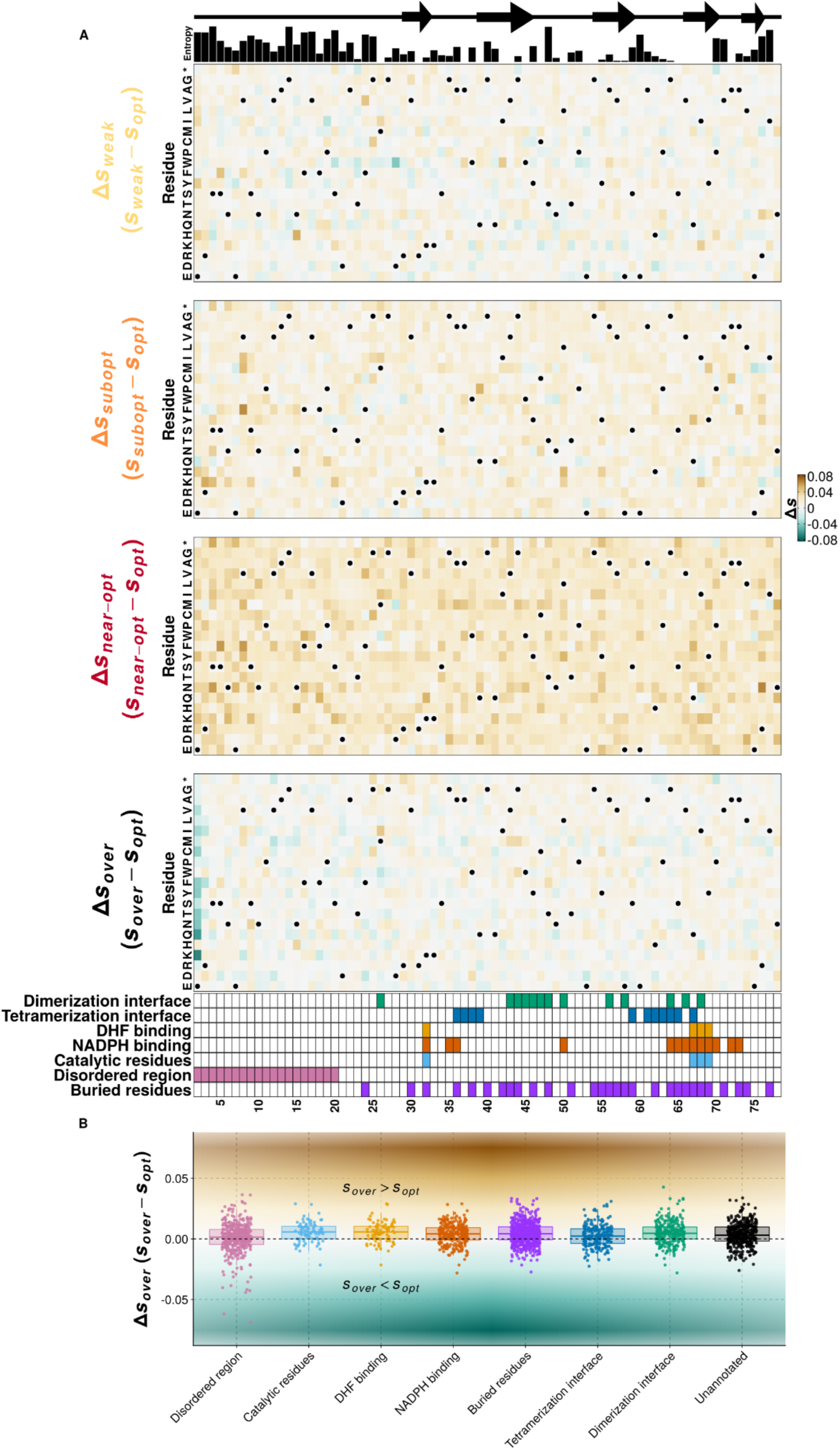
The experiment without selection for DfrB1 activity does not recapitulate the masking effects of higher expression. (**A**) Differences between selection coefficients observed at optimal expression and other expression levels in the experiment without selection for DfrB1 activity (without TMP). WT residues are labeled with dots. (B) Distributions of differences in fitness effects **Δ***s_over_* (*s_over_* - *s_over_*) observed for different features of DfrB1.

**Fig. S24.**
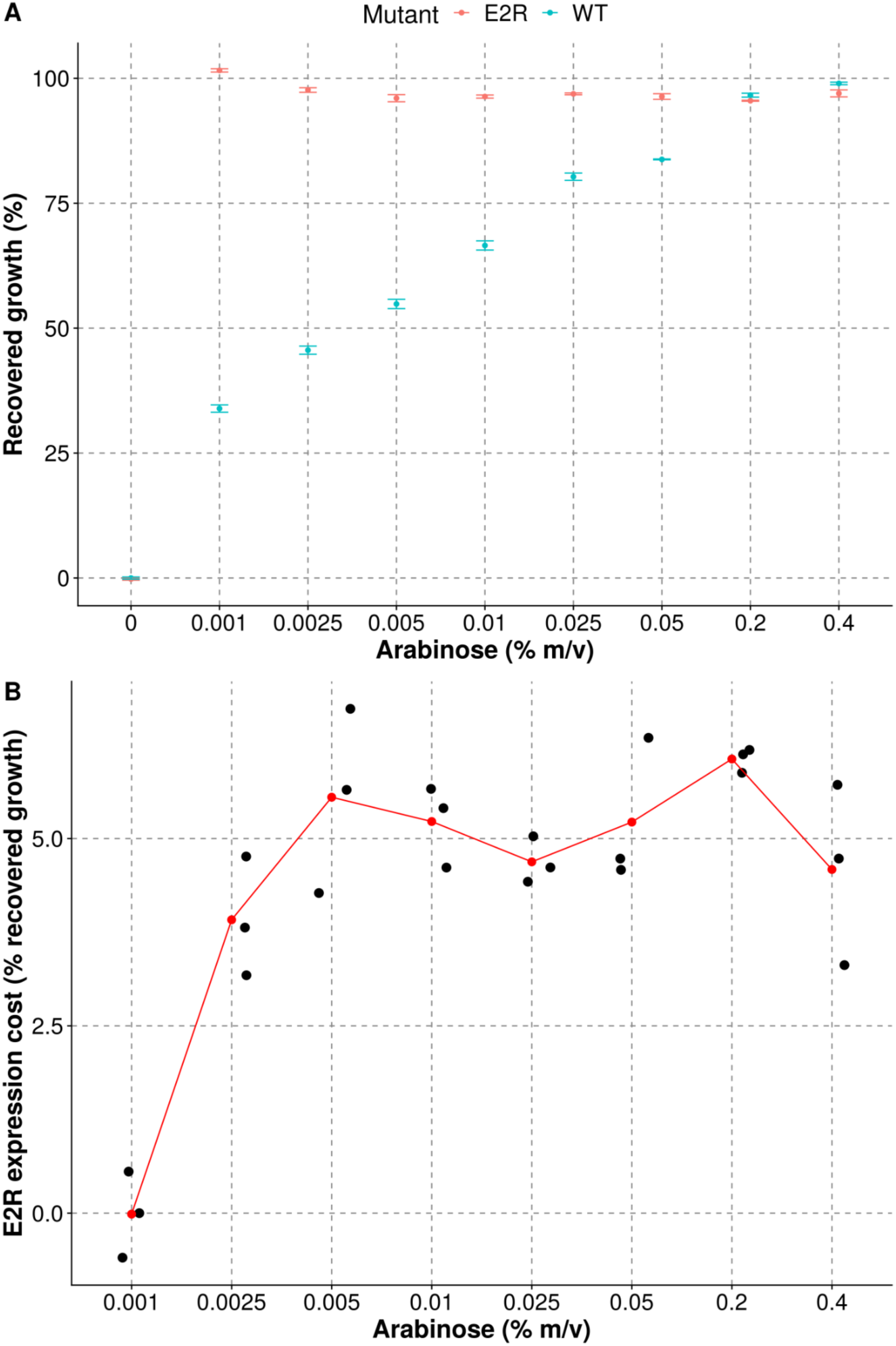
Comparison of growth recovery for WT DfrB1 and E2R mutant at different expression levels. (**A**) Recovered growth is evaluated as the relative difference in growth (area under the curve) of the strain in medium with TMP relative to the same medium without TMP for a given arabinose concentration. Dots and error bars indicate the mean and the standard error of the mean of three replicates. (B) Cost of expression for the E2R mutant in terms of percentage of recovered growth compared to the median recovery at 0.001 % arabinose. The red dots indicate the mean of three replicates for each arabinose concentration.

